# The core root microbiome of *Spartina alterniflora* is predominated by sulfur-oxidizing and sulfate-reducing bacteria in Georgia salt marshes, USA

**DOI:** 10.1101/2021.07.06.451362

**Authors:** Jose L. Rolando, Max Kolton, Tianze Song, J.E. Kostka

## Abstract

**Background:** Salt marshes are dominated by the smooth cordgrass *Spartina alterniflora* on the US Atlantic and Gulf of Mexico coastlines. Although soil microorganisms are well known to mediate important biogeochemical cycles in salt marshes, little is known about the role of root microbiomes in supporting the health and productivity of marsh plant hosts. Leveraging *in situ* gradients in aboveground plant biomass as a natural laboratory, we investigated the relationships between *S. alterniflora* primary productivity, sediment redox potential, and the physiological ecology of bulk sediment, rhizosphere, and root microbial communities at two Georgia barrier islands over two growing seasons.

**Results:** A marked decrease in prokaryotic alpha diversity with high abundance and increased phylogenetic dispersion was found in the *S. alterniflora* root microbiome. Significantly higher rates of enzymatic organic matter decomposition, as well as the relative abundances of putative sulfur (S)-oxidizing, sulfate-reducing, and nitrifying prokaryotes correlated with plant productivity. Moreover, these functional guilds were overrepresented in the *S. alterniflora* rhizosphere and root core microbiomes. Core microbiome bacteria from the *Candidatus* Thiodiazotropha genus, with the metabolic potential to couple S oxidation with C and N fixation, were shown to be highly abundant in the root and rhizosphere of *S. alterniflora*.

**Conclusions:** The *S. alterniflora* root microbiome is dominated by highly active and competitive species taking advantage of available carbon substrates in the oxidized root zone. Two microbially-mediated mechanisms are proposed to stimulate *S. alterniflora* primary productivity: (i.) Enhanced microbial activity replenishes nutrients and terminal electron acceptors in higher biomass stands, and (ii.) coupling of chemolithotrophic S oxidation with carbon (C) and nitrogen (N) fixation by root and rhizosphere associated prokaryotes detoxify sulfide in the root zone while potentially transferring fixed C and N to the host plant.

## 1. Background

Salt marsh ecosystems are structured by intertidal plant communities at the land-sea interface. Salt marshes are mostly distributed outside of the tropics and comprise a global area of ~5.5 Mha, with approximately 30% of its area located in the continental USA (Mcowen et al., 2017). On the US coastlines of the Atlantic Ocean and Gulf of Mexico, salt marshes are dominated by the smooth cordgrass *Spartina alterniflora* (Mitsch and Gosselink, 1993). Salt marsh ecosystems are biogeochemical hotspots characterized by high rates of primary productivity, organic matter mineralization, and nutrient cycling (Howarth, 1984; Kostka et al., 2002a; Kirwan et al., 2009). As a consequence of their high biological activity, *S. alterniflora* dominated salt marshes provide a broad range of ecosystem services to local and global human populations (Barbier et al., 2011). Ecosystem services provided by salt marshes include estuarine water purification, coastal protection from storm surges, sediment erosion control, maintenance of fisheries, carbon sequestration, and much more (Barbier et al., 2011, Hopkinson et al., 2019).

At the local scale, bottom-up control of *S. alterniflora* primary productivity has been associated with nitrogen (N) uptake kinetics (Mendelssohn and Morris, 2000). Low sediment redox potential and high sulfide concentration have been shown to reduce *S. alterniflora*’ root energy status, decreasing the plant’s available energy for N uptake (Morris and Dacey, 1984; Bradley and Morris, 1990; Koch et al., 1990; Mendelssohn and McKee, 1992). Thus, naturally occurring gradients of *S. alterniflora* primary productivity are usually found as a function of the plant’s distance to large tidal creeks (Valiela et al., 1978). Sediments at closer proximity to large tidal creeks are flushed more frequently; supplying oxygen, exchanging porewater nutrients, and oxidizing toxic metabolic products such as sulfide. Conversely, areas in the interior of the marsh tend to be stagnant, accumulating chemically reduced and toxic compounds in their interstitial porewater. The microbial mediation of major biogeochemical cycles along this natural gradient in aboveground *S. alterniflora* biomass has been extensively studied (Kostka et al., 2002a; Koretsky et al., 2003; Dollhopf et al., 2005; Hyun et al., 2007; Tobias and Neubauer, 2019; Murphy et al., 2020). However, the relationship between root-microbial interactions and *S. alterniflora* primary productivity has not been characterized in detail.

Biogeochemical evidence points to tightly coupled interactions between *S. alterniflora* and microbial activity in the root zone, which facilitate the rapid exchange of electron (e^−^) acceptors (O_2_, NO_3_, Fe^3+^) and donors (e.g., rhizodeposits, reduced sulfur compounds). For example, reduced sulfur compounds such as pyrite store high amounts of chemically reduced energy, and their oxidation has long been hypothesized to be an important process limiting the energy flow of salt marsh ecosystems (Howarth, 1984). Part of this energy has been speculated to be used to enhance plant growth (Howarth 1984, Morris et al., 1996), similar to the well-known symbiotic relationship between invertebrates and autotrophic sulfur-oxidizing bacteria in marine ecosystems (Dubilier et al., 2008). However, the *S. alterniflora* root-associated microbiome remains largely unexplored.

Most previous studies investigating the *S. alterniflora* root microbiome focused on understanding the factors influencing the activity and taxonomic diversity of root-associated nitrogen-fixing bacteria or diazotrophs (Whiting et al., 1986; Gandy and Yoch, 1988; Lovell et al., 2000; Brown et al., 2003; Davis et al., 2011). A few reports of other functional guilds have shown that chemolithoautotrophs conserving energy through of S, Fe, and ammonium oxidation are also enriched in the *S. alterniflora* root zone when compared to bulk sediment (Thomas et al., 2014; Zheng et al., 2016; Kolton et al., 2020). A drawback of these previous studies is that the majority did not thoroughly separate the root-associated compartment from the surrounding rhizospheric sediment (e.g., Thomas et al., 2014; Zogg et al., 2018; Kolton et al., 2020). Thus, the ecology of the closely associated *S. alterniflora* root microbiome, and its interaction with the plant host still represents an important knowledge gap to be addressed. In other plants, root microbial communities have been shown to be key players in improving plant resistance to biotic and abiotic stress, outcompeting soil-borne pathogens, modulating plant development, and transferring nutrients for plant uptake (Liu et al., 2019; White et al., 2019; Zhang et al., 2020).

To gain a predictive understanding of the beneficial interactions between a host plant and its associated root microbiome, the ecology and potential physiology of its core microbiome must be investigated. A host’s core microbiome is composed of microbial taxa consistently found associated with host individuals and hypothesized to perform key functions in healthy host-microbiome systems (Shade and Handelsman, 2011). The *S. alterniflora* core root microbiome has yet to be defined. Understanding of *S. alterniflora* root microbiome could pave the way to harnessing plant-microbe interactions for the adaptive management and restoration of salt marsh habitats.

Thus, this study sought to elucidate the plant-microbe interactions linked to *S. alterniflora* primary productivity over the course of 2 years at 2 sites in GA, USA. The objectives of the study were to: 1) Evaluate the taxa that constitute the *S. alterniflora* rhizosphere and root core microbiome in GA, USA 2) Characterize the potential metabolism, physiology, and ecology of the prokaryotic taxa enriched at closer proximities to *S. alterniflora* roots along primary productivity gradients. 3) Propose a mechanistic understanding of the relationship between *S. alterniflora* and its root-associated prokaryotic community.

## 2. Results

### 2.1 Primary productivity gradient

Eight transects along a *S. alterniflora* primary productivity gradient were studied in two barrier islands in the state of Georgia, USA. A total of 24 sampling points were established and sampled during the years 2018 and 2019 in the Georgia Coastal Ecosystem – Long Term Ecological Research (GCE-LTER) site 6 at Sapelo Island (Lat: 31.389° N, Long: 81.277° W), and the Saltmarsh Ecosystem Research Facility (SERF) adjacent to the Skidaway Institute of Oceanography on Skidaway Island (Lat: 31.975° N, Long: 81.030° W) (Fig. S1). *S. alterniflora* shoot height and biomass revealed a pronounced primary productivity gradient with ranges observed within sampled points from 16.5 cm to 128.4 cm, and 1.4 g m^−2^ to 1769.8 g m^−2^, respectively. *S. alterniflora* plants were operationally classified in three phenotypes based on shoot height: short (< 50 cm), medium (50 – 80 cm) and tall (> 80 cm). Shoot biomass averaged 149.4 g m^−2^, 307.8 g m^−2^, and 958.7 g m^−2^ in the short, medium and tall phenotype zones, respectively. Although shoot biomass was not closely associated with changes in leaf N concentration and total inorganic N in interstitial porewater, a strong relationship with leaf δ^15^N and sediment C:N ratio was observed (Fig. 1a, 1b, S2, S3, S4). Higher shoot biomass was also associated with zones in the marsh with elevated sediment redox potential (Eh), which was evidenced not only by direct redox measurements, but also by higher concentrations of interstitial Fe^3+^, and NO3-in the zones dominated by the tall *S. alterniflora* phenotype (Fig. 1c, Fig. S3). Conversely, in zones dominated by the short and medium phenotypes, concentrations of porewater ∑S^2−^ were elevated, reaching up to 1.5 mM (Fig. S3). Shoot biomass also showed a negative relationship with leaf temperature, a proxy for leaf stomatal conductance (Ramírez et al., 2016; Fig. 1d).

**Figure 1:**
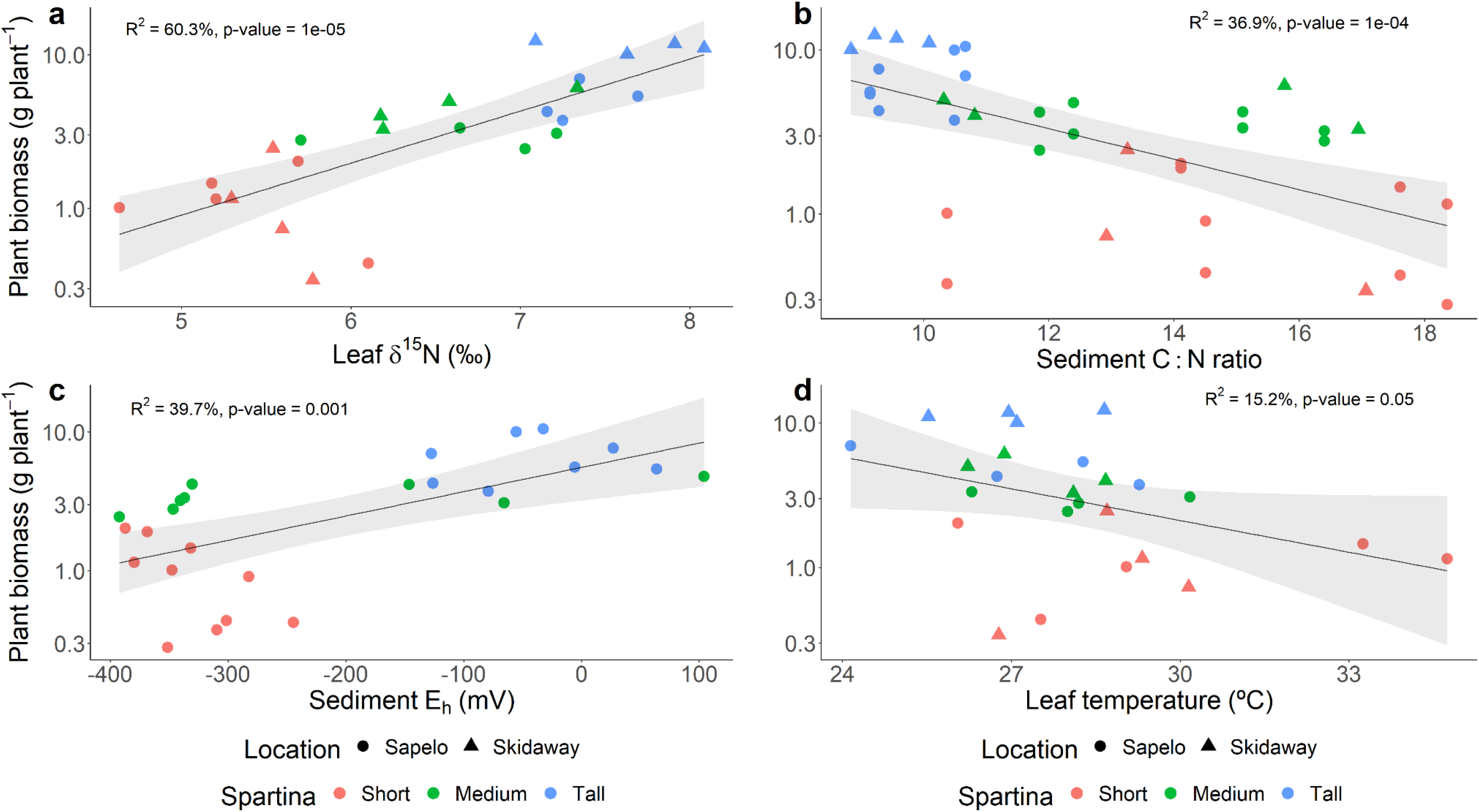
Statistical comparisons by linear regression analysis of average plant biomass and leaf natural ^15^N abundance (δ^15^N) (a), sediment C:N ratio (b), sediment redox potential (Eh) (c), and leaf temperature (d). Plant biomass was calculated as the average of 10 individuals per sampling point. Each leaf δ^15^N, sediment C:N ratio, redox potential, and leaf temperature value represents the average of 3, 1, 3, and 5 replicates, respectively.

Rates of extracellular enzyme activity for enzymes that catalyze the catabolism of organic C, N and P compounds, showed a strong relationship to *S. alterniflora* primary production. Rates of extracellular β-glucosidase (C), chitinase (C, and N) and phosphatase (P) activity in homogenized sediment slurries were consistently higher in zones with greater *S. alterniflora* shoot biomass at both Sapelo and Skidaway Island (Fig. 2).

**Figure 2:**
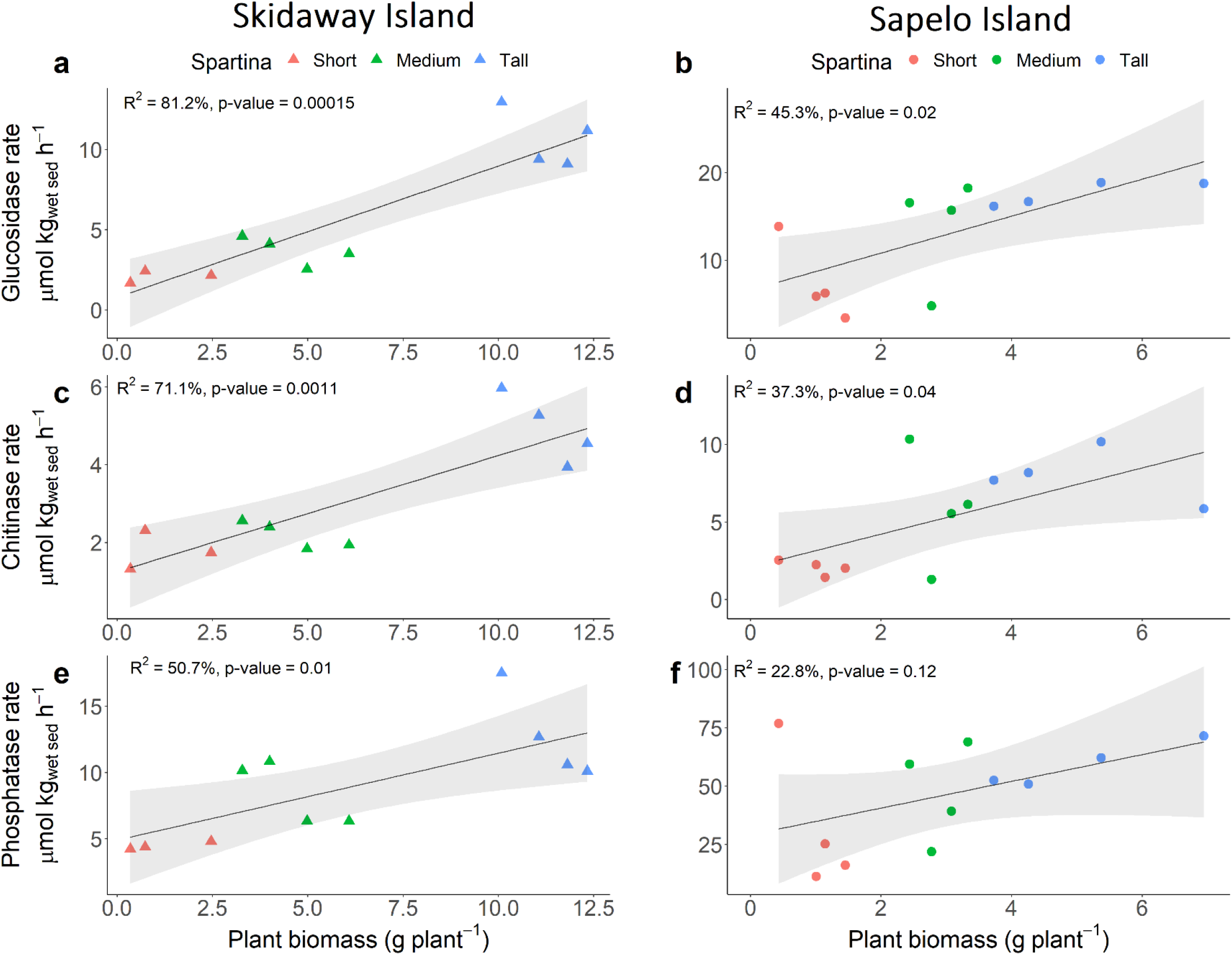
Statistical comparisons by linear regression analysis of enzyme activity rates and plant biomass assessed at Skidaway Island (a), and Sapelo Island (b). For each sample, rates were calculated from 8 time point measurements. Plant biomass represents the average of 10 individuals per sampling point.

### 2.2 Microbiome diversity

Prokaryotic diversity and abundance were investigated across the *S. alterniflora* biomass gradient in three compartments: bulk sediment, rhizosphere, and root. The root compartment was recovered by sonication in an epiphyte removal buffer; thus, likely containing mostly endosphere with residual rhizoplane microbial communities (Simmons et al., 2018). A total of 32,740 unique amplicon sequence variants (ASVs) were inferred using DADA2 v.1.10 (Callahan et al., 2016). After quality filtering, 10,068,980 high quality SSU rRNA sequence reads with a median depth of 49,619 reads per sample were used for microbiome analysis (further details in Materials and Methods). Prokaryotic communities associated with the tall *S. alterniflora* bulk and rhizosphere sediment were more diverse and abundant when compared to those of the short phenotype (Fig. 3a, Fig. 3b). In the root compartment, alpha-diversity and prokaryotic abundance were highest in the short phenotype (Fig. 3a, Fig. 3b). The root compartment showed a significant decline in alpha diversity in all plant phenotypes driven by a decrease in both richness and evenness when compared to their bulk and rhizospheric counterparts (Fig. 3a, Fig. 3c). Mean richness ± 95% confidence interval (CI_95%_) across microbiome compartments was 922 ± 40, 981 ± 33, and 695 ± 40 observed ASVs for bulk sediment, rhizosphere and root prokaryotic communities, respectively. Decreased evenness in the root compartment was evidenced by the presence of highly dominant ASVs (Fig. 3c). Despite the decrease in prokaryotic diversity in the root, abundance remained high, in the range of 10^7^ SSU rRNA gene copies g^−1^_fresh root_ in both tall and short *S. alterniflora* phenotypes (Fig. 3b).

**Figure 3:**
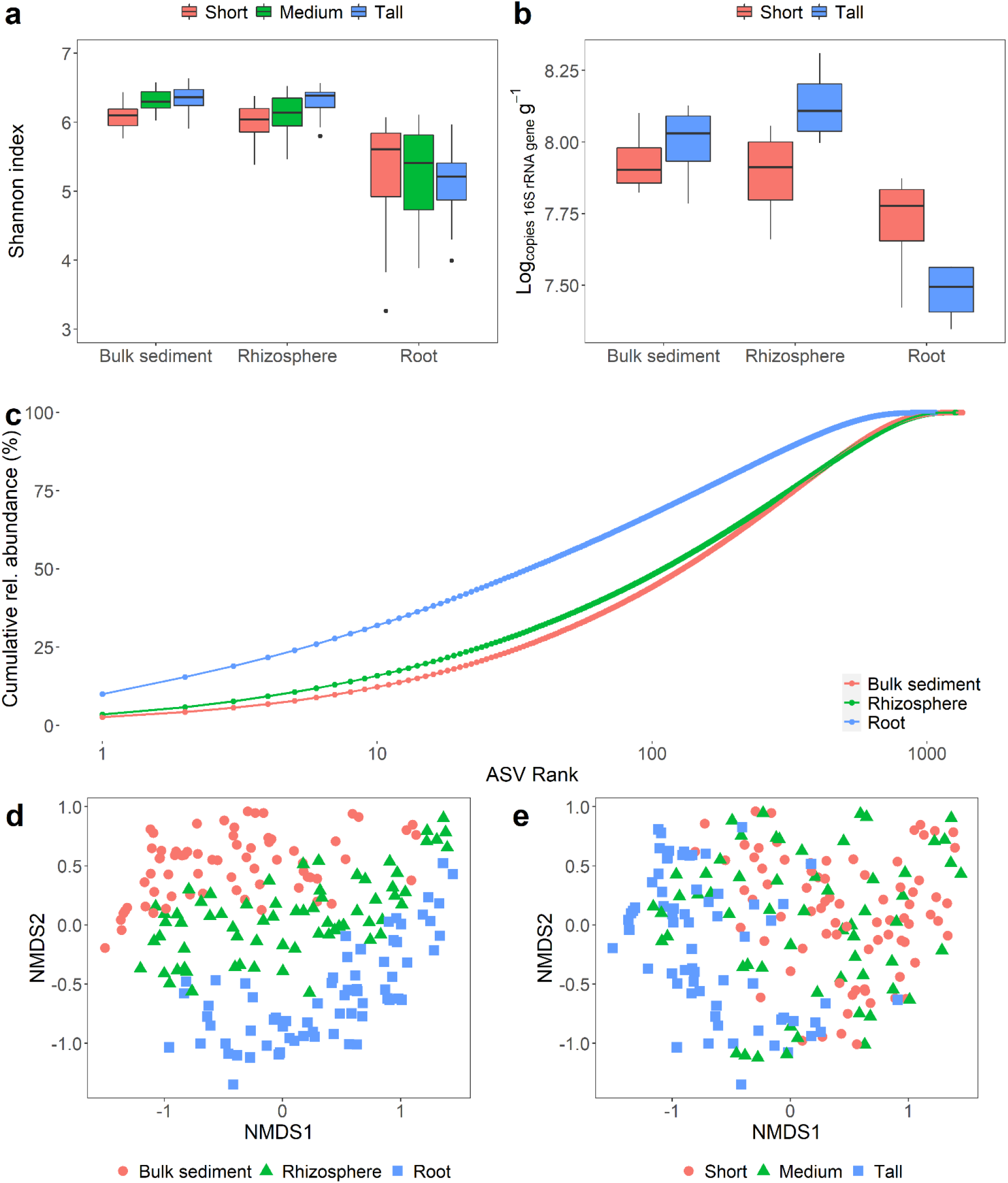
Diversity and abundance of the *S. alterniflora* microbiome. Boxplots of the Shannon diversity index (a), and prokaryotic abundance determined by qPCR of SSU rRNA genes (b) per microbiome compartment and *S. alterniflora* phenotype. Evenness across plant compartments assessed by a cumulative rank-abundance plot (c). Non-metric multidimensional scaling (nMDS) ordination of the Bray-Curtis dissimilarity matrix across all collected samples with colors representing microbiome compartment (d) and *S. alterniflora* phenotype (e). nMDS stress: 0.10.

*S. alterniflora* phenotype and microbiome compartment were the most significant deterministic forces controlling prokaryotic community assembly in GA salt marshes (Fig. 3de, Table 1). PERMANOVA analysis using the Bray-Curtis dissimilarity index showed that *S. alterniflora* phenotype explained less species exchange in the root when compared to the other two compartments (Table 1).

**Table 1:** Analysis of the deterministic parameters controlling microbiome assembly. PERMANOVA analysis was conducted using the Bray-Curtis metric with 999 permutations. Results are provided for the complete data set and for microbiome compartments.

| Factor | All |  | Bulk sediment |  | Rhizosphere |  | Root |  |
| --- | --- | --- | --- | --- | --- | --- | --- | --- |
|  | F-value | R<sup>2</sup> | F-value | R<sup>2</sup> | F-value | R<sup>2</sup> | F-value | R<sup>2</sup> |
| Compartment | 16.6 | 12.1** | - | - | - | - | - | - |
| *S. alterniflora* phenotype | 12.0 | 8.8** | 8.2 | 17.6** | 7.1 | 16.1** | 4.2 | 10.7** |
| Location | 13.9 | 5.1** | 9.2 | 9.8** | 8.1 | 9.2** | 4.7 | 5.9** |
| Depth | 4.6 | 1.7** | 3.2 | 3.5** | 2.4 | 2.7** | 2.7 | 3.4** |
| Year | 2.3 | 0.8** | 2.6 | 2.8** | 1.5 | 1.7 | 1.6 | 2.0* |

Phylogenetic community structure was assessed with the nearest taxon index (NTI) and beta nearest taxon index (βNTI) for within and between prokaryotic communities, respectively (Stegen et al., 2012, 2013). An NTI value greater than 2 indicates greater phylogenetic clustering within a community than expected by chance. All bulk and rhizospheric prokaryotic communities had an NTI greater than 2; while 91% of the root prokaryotic communities met this threshold. NTI values decreased in closer proximity to roots (Fig. S5). Average ± CI_95%_ NTI values per microbiome compartment were 7.0 ± 0.3, 5.6 ± 0.2, and 3.8 ± 0.3 for prokaryotic communities from bulk sediment, rhizosphere and root compartments, respectively. In order to assess phylogenetic turnover between similar environments in the investigated salt marshes, βNTI values between samples from the same microbiome compartment, *S. alterniflora* phenotype, year and location were calculated. An βNTI value lower than −2 indicates less phylogenetic turnover between samples than expected by chance. Pairwise comparison between bulk sediment, rhizosphere, and root prokaryotic communities revealed that 92.2%, 92.6% and 77.8% βNTI values were below −2, respectively. Similar to NTI analysis, a trend of lower phylogenetic relatedness was observed closer to the root (Fig. S5). Average ± CI_95%_ βNTI was −5.2 ± 0.3, −4.6 ± 0.3, and −3.1 ± 0.2 in bulk sediment, rhizosphere and root pairwise comparisons, respectively. The microbiome compartment effect on NTI and βNTI was consistent in all *S. alterniflora* phenotypes.

### 2.3 *S. alterniflora* Root-associated prokaryotic community composition

Overall, at the phylum level, prokaryotic communities were predominated by ASVs from the *Proteobacteria* (46.7%), *Chloroflexi* (15.2%), *Bacteroidetes* (8.4), *Epsilonbacteraeota* (3.7%), *Spirochaetes* (3.6%), and *Acidobacteria* (3.5%) phyla (Fig. S6). At higher *S. alterniflora* biomass, an increase in the relative abundance of *Proteobacteria* ASVs and a decline in the relative abundance of *Chloroflexi* and *Spirochaetes* ASVs was observed (Fig. S6). At increasing proximity from the root, the relative abundance of *Proteobacteria*, *Spirochaetes* and *Epsilonbacteraeota* increased while *Acidobacteria* and *Bacteroidetes* decreased (Fig. S6). Prokaryotic taxa with the potential to catalyze redox reactions in the S, Fe, and N cycles were investigated in greater detail due to their known significance in salt marsh ecosystem functioning. Putative function was inferred based on homology at the genus level with described prokaryotic species (Table S1). Prokaryotes putatively capable of nitrification (a.k.a. nitrifiers) exhibited higher relative abundance in areas colonized by the tall *S. alterniflora* phenotype, in comparison to areas occupied by the short and medium phenotypes (Fig. 4a). Dominant nitrifiers in the studied system included members of the bacterial genera *Candidatus* Nitrotoga and *Nitrospira*, as well as the archaeal genus *Candidatus* Nitrosopumilus (Fig. 4a). Additionally, a significant enrichment in taxa potentially involved in the Fe and S cycles were detected in the plant root relative to the bulk sediment (Fig. 4b,c,d). The putative Fe-oxidizer of the *Zetaproteobacteria*, *Mariprofundus* sp. showed high relative abundance in the roots of the tall *S. alterniflora* phenotype, while *Acidihalobacter* of the *Gammaproteobacteria* was the predominant Fe-oxidizer in the roots of the short phenotype (Fig. 4b). Putative autotrophic endosymbionts capable of S oxidation from the *Candidatus* Thiodiazotropha genus and *Thiomicrospirales* order preferably colonized the roots of *S. alterniflora* phenotypes regardless of plant phenotype (Fig. 4c). Sulfur oxidizers from the S*ulfurovum* genus preferentially colonized the areas dominated by the short *S. alterniflora* phenotype in all compartments (Fig. 4c). Putative sulfate-reducers of the *Desulfobacterales* order: *Desulfatitalea*, *Desulfopila*, and *Desulfosarcina* genera were enriched at closer proximities to the *S. alterniflora* root (Fig. 4d).

**Figure 4:**
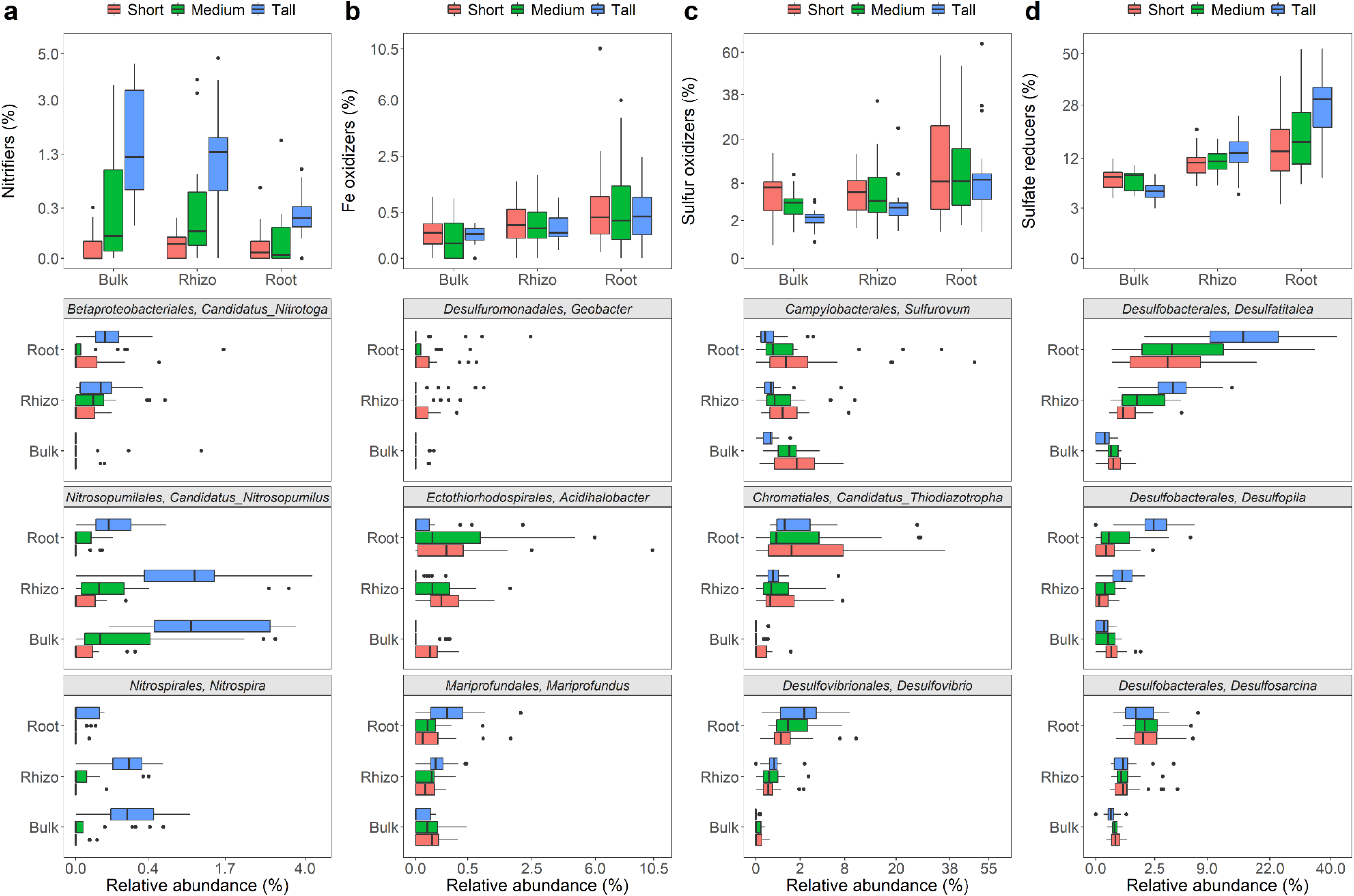
Relative abundance of putative nitrifiers (a), Fe oxidizers (b), S oxidizers (c), and S/sulfate reducers (d) by microbiome compartment and *S. alterniflora* phenotype.

Based on differential abundance analysis performed in DESeq2, many ASVs were shown to be significantly enriched in the root compartment. Interestingly, many of the enriched taxa appear to be capable of N fixation, including putative sulfur oxidizers *Candidatus* Thiodiazotropha, *Desulfovibrio*, and *Arcobacter*, S/sulfate reducers *Sulfurospirillum,* and *Desulfatitalea*, and *Novosphingobium*, *Azoarcus*, and *Celerinatantimonas* bacteria (Fig. S7a). Taxa significantly enriched in the tall *S. alterniflora* phenotype comprised nitrifiers from the *Nitrospira* and *Candidatus* Nitrosopumilus genus, putative metal (Fe and Mn) reducer *Georgfuchsia*, and a diverse set of aerobic or facultative anaerobic chemoheterotrophs (Fig. S7b).

### 2.4 *S. alterniflora* core microbiome

Taxa consistently found in independent host microbiome samples have been suggested to perform key functions in healthy host-microbiome interactions (Shade and Handelsman, 2011), and the set of persistent taxa have been defined as the host’s core microbiome. For this study, an ASV prevalence threshold was operationally defined by plotting the relative abundance and richness of the rhizosphere and root core microbiomes at 10% intervals from 0% to 100% ASV prevalence cutoffs (Fig. S8). A conservative prevalence cutoff of 60% was determined by visually inspecting a threshold in which richness remained stable at increasing cutoff values (Fig. S8). The *S. alterniflora* core root microbiome was composed of only 38 out of 14,505 ASVs, and 54 out of 19,435 ASVs in the root and rhizosphere, respectively. However, in both cases the core microbiome comprised approximately 20% relative abundance of the total prokaryotic community (Fig. S8). Both the root and the rhizosphere core microbiomes were dominated by taxa with inferred metabolic potential for S redox reactions (Fig. 5). The *S. alterniflora* root core microbiome was comprised of putative autotrophic S oxidizers of the *Sulfurovum* and *Candidatus* Thiodiazotropha genera (Fig. 5), while sulfate reducers were represented by ASVs from the *Desulfatiglans*, *Desulfocarbo*, *Desulfatitalea*, *Desulfobulbus*, *Desulfopila*, *Desulfosarcina*, SEEP-SRB1 and Sva0081 genera (Fig. 5). Core root taxa with diazotrophic potential included ASVs from the *Candidatus* Thiodiazotropha, *Desulfatitalea*, *Desulfobulbus* and *Spirochaeta* genera (Fig. 5). In the rhizosphere, the proportion of ASVs with unknown classification at the genus level according to the SILVA database (release 132), comprised up to ~60% relative abundance of the core microbiome. Core rhizosphere ASVs with the putative capacity for S oxidation included members of the *Sulfurovum* and *Thioalkalispira* genera (Fig. 5). Similar to the root core microbiome, nearly half of the identified taxa at the genus level in the core rhizosphere presented sulfate reduction capability, such as ASVs from the *Desulfatiglans*, *Desulfocarbo*, *Desulfatitalea*, *Desulfosarcina*, SEEP-SRB1, and Sva0081 genera (Fig. 5). Putative nitrifiers from the *Candidatus* Nitrosopumilus genus were members of the rhizosphere core microbiome (Fig. 5). Finally, taxa with N fixation capability in the rhizosphere core microbiome included putative sulfur oxidizers from the *Thioalkalispira* genus, sulfate reducers from the *Desulfatitalea* genus, and bacterium from the *Spirochaeta* genus (Fig. 5).

**Figure 5:**
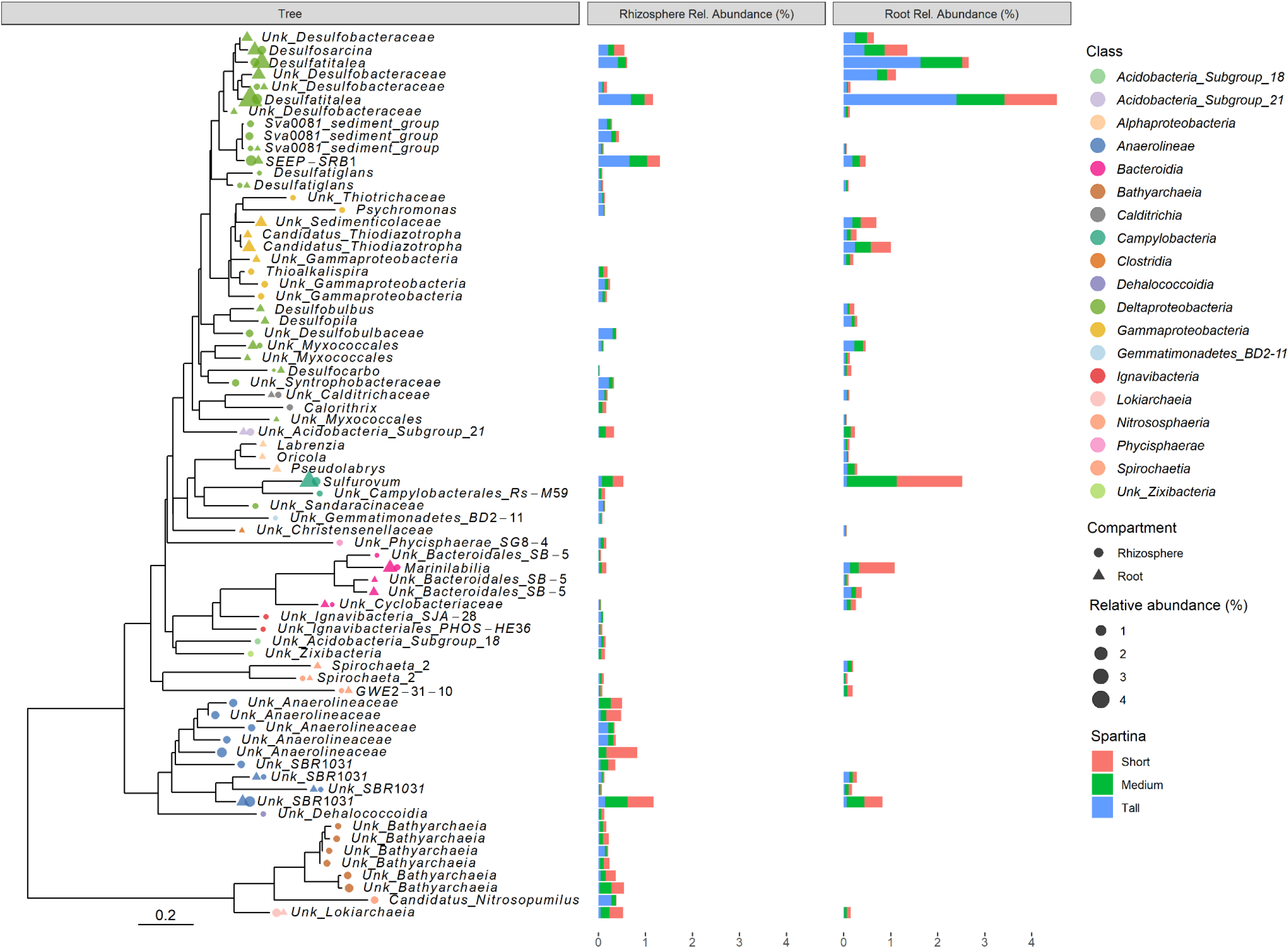
Prokaryotic identity and relative abundance of the *S. alterniflora* rhizosphere and root core microbiome. Phylogenetic characterization was conducted using an approximately-maximum-likelihood model of the 72 ASVs comprising the *S. alterniflora* core rhizosphere and root microbiome. Taxonomic information at the genus level is provided for all ASVs. When taxonomic assignment was unknown at the genus level, the “Unk” prefix was used before the highest resolution taxonomic level assigned.

## 3. Discussion

### 3.1 Biogeochemical processes linked to *S. alterniflora* primary productivity at the local scale

*S. alterniflora* primary productivity is strongly linked to N uptake kinetics in field and lab studies (Mendelssohn et al., 1979; Howes et al., 1986, Mendelssohn and Morris, 2000). At the local scale, reduced, anoxic, and sulfidic root conditions were shown to lead to a decline in root energy status, affecting ammonium uptake kinetics (Koch et al., 1990; Mendelssohn and Morris, 2000). In parallel, elevated salinity has been associated with a decrease in stomatal conductance and photosynthetic activity, along with an increase in dark respiration (Giurgevich and Dunn, 1979; Hwang and Morris, 1994). Our observations corroborate these past results showing that *S. alterniflora* primary productivity is hampered at reduced sediment Eh and under highly sulfidic conditions, when the plants experience limited leaf gas exchange. Results from this study also support our previous research which revealed the dynamic interplay between the growth/physiology of macrophyte plants and macrofaunal bioturbation (Kostka et al., 2002a; Gribsholt et al., 2003), with crab burrow density shown to directly correlate with aboveground plant biomass and sediment redox potential (Fig S2).

Significant differences in stable N isotope composition of sediment and *S. alterniflora* leaves provide evidence that N sources and dynamics are distinct along the studied *S. alterniflora* primary productivity gradient (Craine et al., 2015). We argue that elevated leaf and sediment δ^15^N in the tall *S. alterniflora* phenotype is the result of (i) greater nitrogen loss by coupled nitrification-denitrification, and (ii) differences in the source of N input. Denitrification in Georgia salt marshes is limited and tightly coupled to prokaryotic nitrification, which discriminates against ^15^N (Dollhopf et al., 2005, Craine et al., 2015). Prokaryotes capable of nitrification show higher relative abundance in sediments dominated by the tall *S. alterniflora* phenotype (Fig. 4a), and their activity is known to be inhibited by sulfide toxicity (Joye and Hollibaugh, 1995; Dollhopf et al., 2005). Porewater sulfide concentration in the short and medium *S. alterniflora* phenotype was an order of magnitude higher than in the tall phenotype. A second explanation for δ^15^N enrichment in the tall *S. alterniflora* phenotype is the greater source of planktonic N at closer proximities to the tidal creek. Planktonic tissue has a δ^15^N signature of 8.6 ± 1.0 ‰ (Peterson, 1999), which is more similar to the leaf δ^15^N measured in the tall *S. alterniflora* (7.6 ± 0.3 ‰) when compared to the short phenotype (5.5 ± 0.4 ‰) (Fig. S4). A larger discrepancy in δ^13^C between sediment and leaf tissue in the tall *S. alterniflora* phenotype, as well as a lower sediment C:N ratio, support our interpretation of a reduced relative contribution of vascular plant material to organic matter diagenesis (Fig. S4; Ember et al., 1987; Fig. 1b, Gebrehiwet et al., 2008). The average ± CI_95%_ δ^13^C signature in sediments from the tall *S. alterniflora* phenotype (−20.3 ± 0.4 ‰) more closely resembles that of reported phytoplankton, which enters the marsh via tidal creeks (−17 to −24‰, Fogel et al., 1989; —21.3 ± 1.1 ‰, Peterson, 1999; −20.16 ‰, Dai et al., 2005). The C:N ratio is considered as a proxy for soil organic matter reactivity, and a lower C:N ratio indicates a greater potential for rapid biodegradation and N mineralization (Janssen, 1996). In agreement with this interpretation, sediments with low C:N ratios from the tall *S. alterniflora* zone contained higher prokaryotic biomass, and higher rates of extracellular enzyme activities involved in the C, N, and P cycles. We propose that plant primary productivity is enhanced in the tall *S. alterniflora* zone in part due to more rapid microbial mineralization of higher-quality sediment organic matter of planktonic origin, with released inorganic nutrients then made available for plant uptake.

### 3.2 Assembly of the *S. alterniflora* root microbiome

Previous studies have reported contradictory results with regard to the relationship between the diversity of the *S. alterniflora* microbiome, plant productivity, and proximity to the root (Zogg et al., 2018; Lin et al., 2019; Kolton et al., 2020). Our finding of higher prokaryotic alpha diversity in bulk and rhizospheric sediment associated with the tall *S. alterniflora* phenotype is consistent with previous findings from Skidaway Island, GA (Kolton et al. 2020). However, Zogg et al. (2018) and Lin et al., (2019) did not observe significant differences in prokaryotic alpha diversity between the tall and short *S. alterniflora* phenotypes in bulk and rhizospheric sediments from New England and Guangdong province in China, respectively. Contrasting results could be due to limitations in methodology and experimental design in previous work. Sequencing platforms continue to evolve, enabling higher sequence coverage at lower cost, and our sampling effort was approximately an order of magnitude more intensive than previous studies characterizing the *S. alterniflora* root microbiome, including our own past work (Zogg et al., 2018; Lin et al., 2019; Kolton et al., 2020). Moreover, recently developed *in silico* technology to infer ASVs instead of clustering sequences into operational taxonomic units (OTUs) could also impact the estimation of alpha diversity metrics.

Few studies have investigated microbial diversity in the roots of wetland plant roots. Nonetheless, consistent with our results, a single report from *S. alterniflora* and studies from other wetland/estuarine plants show a decline in diversity in the root compared to the bulk and rhizosphere compartments (Edwards et al., 2015; Hong et al., 2015; Martin et al., 2020a). Prokaryotic abundance in the endosphere has been previously estimated in the 10^4^ to 10^8^ cells g^−1^ range across an array of plant species (Bulgarelli et al., 2013). Taking into consideration that many of these previous estimates employed cultivation-based methods, and prokaryotic genomes often contain multiple SSU rRNA operons, our observation of prokaryotic abundance (10^7^ SSU rRNA gene copies g^−1^) would be placed at the high-end of that range. A marked decrease in prokaryotic alpha diversity with high abundances is an indication that the *S. alterniflora* root compartment is enriched in dominant and highly active species taking advantage of labile carbon sources, reduced inorganic compounds, and the oxidized environment found in the *S. alterniflora* root (Carlson and Forrest, 1982; Maricle and Lee, 2002). Increased relative abundances of S and Fe chemolithotrophs, and aerobic and facultative anaerobic chemoorganotrophs at closer proximities to the root further support this interpretation.

Microbial community assembly in the root endosphere has been proposed as a two-step colonization process (Bulgarelli et al., 2013). The first step is driven by microbial proliferation in the rhizosphere by species capable of utilizing plant-released substrates; while the second is a fine-tuning step in the rhizoplane, where selection by the plant’s genotype-dependent immune system takes place (Bulgarelli et al., 2013). Endospheric microbial species have co-evolved to evade the plant immune system by secreting effector proteins that mimic plant proteins (Trivedi et al., 2020). In *S. alterniflora*, a decrease in prokaryotic richness in the root compartment, and the fact that plant phenotype was the most important deterministic factor assembling the root prokaryotic community, suggest that plant selection is an important process in community assembly. However, community assembly is a result of co-occurring deterministic and stochastic processes (Stegen et al., 2013; Dini-Andreote et al., 2015). Generally, environmental filtering has been shown to be the main deterministic process assembling microbial communities spatially, as evidenced by phylogenetic clustering (Stegen et al., 2012; Freedman and Zak, 2015). Nevertheless, in our study, environmental filtering was relaxed at closer proximity to the root, a microenvironment characterized by an abundance of high-quality e^−^ donors and acceptors. Most likely, increased competition, the dominance of fast-growing bacteria filling a resource-rich niche, and historical contingency (i.e., first prokaryotic species to colonize the root successfully outcompete other taxa) are co-occurring ecological processes reducing the relative importance of environmental filtering in the root zone (Fukami, 2015; Hassani et al., 2018; Toju et al., 2018). Increased species dominance, decreased richness, and increased phylogenetic dispersion with high prokaryotic abundances in the *S. alterniflora* root supports this hypothesis.

### 3.3 Characterization of the *S. alterniflora* core microbiome and potential plant-microbe interactions driving primary productivity

Our results indicate that putative sulfate-reducing and sulfur-oxidizing prokaryotes comprise a large proportion of the *S. alterniflora* root and rhizosphere core microbiomes. Sulfate-reducing communities are comprised of metabolically versatile populations capable of utilizing a broad range of C substrates, including plant-derived substrates (Bahr et al., 2005). The most dominant sulfate-reducer genus in the *S. alterniflora* core root and rhizosphere, *Desulfatitalea*, mainly utilizes short-chain fatty acids as an electron donor and C source (Higashioka et al., 2013). However, other sulfate-reducing members of the root core microbiome such as *Desulfatiglans* and *Desulfocarbo*, have been shown to oxidize plant-derived aromatic compounds (e.g., lignin) (An and Picardal, 2014; Suzuki et al., 2014). In salt marsh ecosystems, biogeochemical data indicates that the C and S cycles are tightly coupled, with *S. alterniflora* photosynthetic activity fueling the activity and C utilization of sulfate-reducers in the rhizosphere (Kostka et al., 2002b; Hyun et al., 2007; Spivak and Reeve, 2015). Similarly, sulfide-oxidizers thrive in the *S. alterniflora* root zone, especially in the short phenotype, where sulfide concentration is elevated (Thomas et al., 2014; Kolton et al., 2020; this study). In our study, ASVs from the *Sulfurovum* genus were enriched in all compartments of the short *S. alterniflora* phenotype. This genus has been described as highly versatile and diverse, allowing for efficient niche partitioning in highly dynamic sulfidic and oxic environments (Meier et al., 2017; Moulana et al., 2020). The co-occurrence of prokaryotic ASVs associated with S anaerobic and aerobic metabolisms in the *S. alterniflora* rhizosphere and root, and highly dynamic O_2_ concentrations at the microscale (Koop-Jakobsen et al., 2018), support the interpretation of a rapid and cryptic S cycle at close proximity to, or even inside, the root tissue. Rapid and coupled cycling of C and S are crucial to the replenishment of nutrients and electron acceptors in the *S. alterniflora* root-zone to support high microbial and plant activity. Rapid organic matter mineralization is especially relevant to effectively recycle N, most often the limiting nutrient for plant productivity in the salt marsh (subsection 3.1).

Previous studies in marshes of the southeastern US show that plant photosynthetic activity and rhizodeposition stimulate N fixation by root-associated sulfate reducers (Whiting et al., 1986; Gandy and Yoch, 1988; Lovell et al., 2000). Consistently, prokaryotic species from the *Desulfobulbus, Desulfatitalea*, *Desulfovibrio,* and *Sulfurospirillum* genera, either significantly enriched or members of the *S. alterniflora* root core microbiome, have been shown to couple sulfate or sulfur respiration to N fixation (Higashioka et al., 2015; Thajudeen et al., 2017). Species from the root and rhizosphere core microbiome genera *Candidatus* Thiodiazotropha and *Thioalkalispira* have the metabolic potential to couple S oxidation with C and N fixation (Barbieri et al., 2010; Petersen et al., 2016). Intriguingly, *Candidatus* Thiodiazotropha is a recently described genus of endosymbionts discovered in the gills of lucinid bivalves (Petersen et al., 2016). Lucinid bivalves are not generally found in salt marshes, but they have been associated with reduced plant sulfide stress in seagrass meadows and mangroves (Lim et al., 2019a; Lim et al., 2019b; Gagnon et al., 2020). In these ecosystems, a tripartite symbiotic relationship occurs, whereby bivalves provide O_2_ to endosymbiont sulfide-oxidizers that detoxify the plant’s environment from sulfide, a known phytotoxin (Gagnon et al., 2020). Recent studies revealed that *Candidatus* Thiodiazotropha is also an important and prevalent member of the seagrass root microbiome, suggesting that these chemolithoautotrophic S oxidizers form a direct association with subtidal marine plant species without the need of a lucinid bivalve partner (Crump et al., 2018; Martin et al., 2020b). Our study confirms this finding, and expands the distribution range of *Candidatus* Thiodiazotropha to the roots of intertidal, estuarine plant species.

Although sulfide is generally considered as a potent phytotoxin, slightly sulfidic conditions (< 1 mM) have been shown to actually stimulate *S. alterniflora* growth in a controlled laboratory experiment (Morris et al., 1996). Energy conservation from sulfide oxidation in the root tissue was speculated to be the driver of increased plant primary production (Mendelssohn and Morris, 2000). Further, sulfide oxidation to sulfate has been demonstrated inside *S. alterniflora* root tissues using isotope tracers (Carlson and Forrest, 1982; Lee et al., 1999). However, it is still not clear what process, biological or chemical, dominates sulfide oxidation inside *S. alterniflora* roots. We propose that *S. alterniflora* shares a symbiotic relationship with S oxidizers in both the rhizosphere and root compartments. Sulfur oxidation may be mediated not only by *Candidatus* Thiodiazotropha bacteria, but also members of the *Sulfurovum* and *Thioalkalispira* genera, or endosymbionts from the *Thiomicrospirales* order. Previously studied microbial species from the *Sulfurovum* genus and endosymbionts from the *Thiomicrospirales* order have been demonstrated to fix C; whereas members from *Desulfovibrio*, *Thioalkalispira*, and *Candidatus* Thiodiazotropha genera have been shown to perform both C and N fixation (Barbieri et al., 2010; Petersen et al., 2016; Suzuki et al., 2006; Thajudeen et al., 2017). Moreover, Crump et al. (2018) studying the root microbiome of seagrass *Zostera* spp., found high transcripts levels of N fixing and sulfur oxidizing genes from *Gammaproteobacteria* species, including endosymbionts of marine invertebrate from the *Sedimenticolaceae* family, which includes the *Candidatus* Thiodiazotropha genus. Given that the *S. alterniflora* root zone is enriched in reduced S and its growth is limited by N uptake, we suggest that diazotrophy coupled to sulfide oxidation may be a key process that was previously overlooked. However, direct measurements of N and C fixation, and sulfur oxidation in the roots of *S. alterniflora*, their rates and controls, along with their relative contribution to plant growth remain unclear and require further research. Given that most studies, including this study, have inferred the coupling of S oxidation and N fixation based on gene homology and/or taxonomic placement, this interpretation should be treated with caution.

## 4. Conclusions

We studied a gradient in *S. alterniflora* productivity to characterize the ecology and physiology of the *S. alterniflora* root-associated microbiome, and its potential role shaping plant physiological performance. In sediments from the tall *S. alterniflora* phenotype, higher prokaryotic biomass and more rapid microbial mineralization of organic matter was linked to greater inorganic nutrients replenishment for plant uptake. Prokaryotic communities from bulk and rhizospheric sediment associated with the tall *S. alterniflora* phenotype contained the highest alpha diversity; while a decline in diversity was observed in the root in comparison to the to bulk and rhizosphere sediment compartments in all *S. alterniflora* phenotypes. A marked decrease in prokaryotic alpha diversity with high abundances and increased phylogenetic dispersion was observed in the *S. alterniflora* root compartment. Thus, we propose that the *S. alterniflora* root microbiome is dominated by highly active and competitive species taking advantage of available carbon substrates in the oxidized root zone. The high relative abundance of prokaryotic ASVs with putative S oxidation and sulfate reduction capability in the *S. alterniflora* rhizosphere and root suggests a rapid S cycle at close proximity to, or even inside, the root tissue. Moreover, both functional guilds were overrepresented in the *S. alterniflora* rhizosphere and root core microbiome. Rapid recycling of S is crucial for organic matter mineralization in anoxic marsh sediments. Thus, we propose that *S. alterniflora* shares a symbiotic relationship with S oxidizing bacteria in both the rhizosphere and root compartments. Sulfur oxidizers may benefit *S. alterniflora* plants not only by removing potentially toxic sulfide from the root zone, but also by coupling S oxidation with N and/or C fixation. The contribution to plant growth of each of these microbial processes represents a knowledge gap that warrants further research.

## 5. Materials and methods

### 5.1 Sampling design and general site description

The study was carried out in two salt marshes for which long-term data is available in the state of Georgia, USA: (i) the Georgia Coastal Ecosystem – Long Term Ecological Research (GCE-LTER) site 6 at Sapelo Island (Lat: 31.389° N, Long: 81.277° W), and (ii) the Saltmarsh Ecosystem Research Facility (SERF) adjacent to the Skidaway Institute of Oceanography on Skidaway Island (Lat: 31.975° N, Long: 81.030° W) (Fig. S1). The GCE-LTER site was sampled twice, during July 2018 and 2019, while the SERF site was sampled once in July 2019. Four ~100 m transects adjacent to large tidal creeks with two to four sampling points along *S. alterniflora* primary productivity gradients were sampled at each site (total: 24 sampling points, Fig. S1).

At each sampling point along the transects, a 50 cm × 50 cm quadrat was established to measure the density of marsh periwinkle snails (*Littoraria irrorata*), and fiddler crab (*Uca pugnax*) burrows. *In-situ* sediment pH and redox potential (Eh) were measured in triplicate at two sediment depths (2.5 and 7.5 cm) with a hand-held pH/ORP meter during the two sampling events at Sapelo Island (HI-98121 tester, Hanna instruments, Woonsocket, RI, USA).

### 5.2 *S. alterniflora* ecophysiology and elemental analysis

*S. alterniflora* shoot density was quantified at every sampling point in 50 cm × 50 cm quadrats, and shoot height measured for 10 plants per sampling point. *S. alterniflora* plants were operationally classified in three phenotypes based on shoot height: short (< 50 cm), medium (50 – 80 cm) and tall (> 80 cm). *S. alterniflora* shoot biomass was estimated by allometry, with an equation calibrated at Sapelo Island (Table S2, Wieski and Pennings, 2014). During the 2019 sampling events, leaf C and N concentration and ^13^C and ^15^N isotopic natural abundance were determined for 3 plants per sampling point with elemental and isotope analyses conducted at the University of Georgia – Center for Applied Isotope Studies (CAIS https://cais.uga.edu/). Leaf elemental analysis was performed by the micro-Dumas method, while isotopic natural abundance was measured by isotope ratio mass spectrometry. ^13^C natural abundance was expressed as the per mille (‰) deviation from the Pee Dee Belemnite standard (PDB) ^13^C:^12^C ratio (δ^13^C); while ^15^N natural abundance expressed as the ‰ deviation from the N2 atmospheric ^15^N:^14^N ratio (δ^15^N). Leaf temperature, a proxy for stomatal conductance (Ramírez et al., 2016), was measured between 11:00 and 12:00 utilizing a Fluke-62 MAX+ infrared thermometer (Fluke Co. USA, Everett, WA). All leaf ecophysiological measurements were performed in young, expanded, and sun exposed leaves.

### 5.3 Porewater and sediment sampling and chemical analysis

Rhizon samplers with 0.15 μm pore size filters (model CSS, Rhizosphere Research Products, Wageningen, The Netherlands, https://www.rhizosphere.com/rhizons), were used to extract sediment porewater at 2.5 and 7.5 cm depth from every sampling point. For each porewater sample, a 2 ml subsample was frozen at −20°C for salinity, nitrate, ammonium, and phosphate concentration analysis; a 2 ml subsample was immediately acidified with 20 μL 12N HCl for Fe^2+^ and Fe^3+^ concentration analysis; and a 100 μL subsample was immediately fixed into 1 ml zinc acetate 2% (w/v) solution for sulfide concentration analysis (Kostka et al., 2002a; Hyun et al., 2007).

Porewater concentrations of nitrate, ammonium, phosphate, sulfide, Fe^3+^, and Fe^2+^ were quantified based on spectrophotometric methods as previously described (Watanabe and Olsen 1965, Cline 1969, Stookey 1970; Sims et al. 1995; Garcia-Robledo et al. 2014). Porewater chloride concentration was determined by HPLC with ultraviolet detection as described by Beckler et al. (2014). Salinity was calculated based on porewater chloride concentration.

Sediment samples were collected at all sampling points from two depth intervals, 0-5 cm and 5-10 cm. An approximately 30 g sediment subsample was oven-dried at 60°C for 72 hours. The oven-dried sample was homogenized using a PowerGen high throughput homogenizer (Fisherbrand, Pittsburgh, PA) and sent to the University of Georgia CAIS (https://cais.uga.edu/) for organic C and N concentration and ^13^C and ^15^N isotopic natural abundance analyses. Sediment organic C and ^13^C isotopic natural abundance was analyzed in acid-fumigated samples (Ramnarine et al., 2011).

Samples collected for microbial community analysis were flash-frozen in an ethanol and dry ice bath, and stored at −80°C until nucleic acid extraction. Sediment samples collected for enzymatic rates were kept at 4 °C, and analyzed within 4 hours after sampling.

### 5.4 Extracellular enzyme activity

Rates of extracellular enzymatic activity (β-glucosidase, phosphatase, and chitinase) were measured in 2019 in Sapelo and Skidaway Island. Technical duplicates of 0-10 cm deep sediment samples were collected for each sampling point. Rates were analyzed in a homogenized sediment slurry using fluorescent 4-methylumbelliferone (MUF) linked-substrates (Table S3). Slurry preparation consisted of mixing wet sediment with 50 mM Tris buffer (pH 7) in a 1:2 (w/v) ratio. The slurry was homogenized with a stomacher homogenizer (model 400; Seward Medical, London, England) at 200 rpm for 30 s in ~1-mm filter bags. Sediment slurry (800 μL) was incubated in the dark with 200 μL of fluorescent-linked substrate (initial substrate concentration: 40 μM). Product accumulation was measured at 0, 0.5, 1, 1.5, 3, 4, 6 and 8 hours after start of the incubation based on fluorescence intensity (excitation: 355 nm, emission: 460 nm) using a microplate fluorescence reader (SpectraMax M2, Molecular Devices, San Jose, CA). Enzymatic rates were calculated by fitting a linear regression, and reported in μmol kg_wet sediment_^−1^ h^−1^.

### 5.5 Plant sampling, compartment fractionation, and molecular biology

*S. alterniflora* roots were sampled at two depths (0-5 and 5-10 cm). Sediment loosely attached to the root was immediately washed two times with creek water in the field. Roots with remaining rhizospheric sediment were flash frozen in an ethanol-dry ice bath and stored at −80°C until analysis.

#### 5.5.1 Plant compartment separation

Separation of the root and rhizosphere compartments was performed by sonication in an epiphyte removal buffer (0.1% (v/v) Triton X-100 in 50 mM potassium-phosphate buffer) (Simmons et al. 2018). Sonication was performed at 4°C for 10 minutes with pulses of 160W for 30 s interspersed with 30 s pauses. Sediment detached from the roots was centrifuged and considered to be the rhizosphere compartment, while the sonicated roots were considered as the root compartment. The root compartment was washed with PBS buffer (pH 7.4) three consecutive times, and ground with liquid N before DNA extraction. Simmons et al. (2018) protocol was designed to isolate endospheric DNA; however, since we did confirm the separation by microscopy, it is possible that our root compartment contained residues of the rhizoplane compartment.

#### 5.5.2 Removal of extracellular DNA

Prior to bulk sediment DNA extraction, extracellular dissolved or sediment-adsorbed DNA was removed according to Lever et al. (2015). Briefly, 2 g of sediment was incubated with 2 ml carbonate dissolution solution (0.43 M sodium acetate, 0.43 M acetic acid, 10 mM EDTA, 100 mM sodium metaphosphate, 3% (w/v) NaCl, pH 4.7) for 1 hour while orbital shaking at 300 rpm and room temperature. Afterwards, 16 ml of a 300mM Tris-HCl and 10mM EDTA solution (3% NaCl, pH 10.0) was added into the slurry and incubated for 1 additional hour at the same orbital shaking condition. After centrifugation at 10,000 g for 20 minutes at room temperature, the pellet was composed by extracellular-DNA free sediment and used for prokaryotic community characterization.

#### 5.5.3 DNA extraction and sequencing library preparation

DNA extraction for all assessed compartments was performed using the DNeasy PowerSoil kit (Qiagen, Valencia, CA) according to the manufacturer’s protocol. The concentration of extracted DNA was determined with the Qubit HS assay (Invitrogen, Carlsbad, CA). Amplification of the SSU rRNA gene V4 region was performed using the primers 515F (5’- GTGCCAGCMGCCGCGGTAA’) and 806R (5’-GGACTACHVGGGTWTCTAAT’) (Caporaso et al., 2011). Reactions were performed in triplicate of 5 ng DNA template in a solution containing DreamTaq buffer, 0.2 mM dNTPs, 0.5 μM of each primer, 0.75 μM of each mitochondrial (mPNA) and plastid (pPNA) peptide nucleic acid (PNA) clamps, and 1.25 U DreamTaq DNA polymerase as previously described (Kolton et al 2020, further details in Table S4). PNA clamps have been shown to reduce plant plastid and mitochondrial DNA amplification in PCR reactions (Lundberg et al., 2013). Triplicate PCR products were pooled together, barcoded with 10-base unique barcodes (Fluidigm Corporation, San Francisco, CA), and sequenced on an Illumina MiSeq2000 platform using a 500-cycle v2 sequencing kit (250 paired-end reads) at the Research Resources Center in the University of Illinois at Chicago. The raw SSU rRNA gene amplicon sequences have been deposited in the BioProject database (http://ncbi.nlm.nih.gov/bioproject) under accessions PRJNA666636.

### 5.6 Quantification of prokaryotic abundance

Prokaryotic abundance was quantified by quantitative polymerase chain reaction (qPCR) of the SSU rRNA gene with general primers in a subset of 24 samples collected in Sapelo Island in 2018 and 2019. The subset comprised superficial (0-5 cm) samples from all three compartments, collected from the four established transects in Sapelo Island. Only sample from the tall and short extreme *S. alterniflora* phenotypes were included into the analysis. Samples were analyzed in triplicate using the StepOnePlus platform (Applied Biosystems, Foster City, CA, United States) and PowerUp SYBR Green Master Mix (Applied Biosystems, Foster City, CA, United States). Reactions were performed in a final volume of 20μl using the standard primer set 515F (5’- GTGCCAGCMGCCGCGGTAA’) and 806R (5’-GGACTACHVGGGTWTCTAAT’) specific for the prokaryotic SSU rRNA gene (Caporaso et al., 2011, Table S4). To avoid plant plastid and mitochondrial DNA amplification from rhizosphere and root samples, peptide nucleic acid PCR blockers (PNA clamps, 0.75 μM) were added to all qPCR reactions (Lundberg et al., 2013). Standard calibration was performed from a 10-fold serial dilution (10^3^ to 10^8^ molecules) of standard pGEM-T Easy plasmids (Promega, Madison, WI, United States) containing target sequences from *Escherichia coli* K12. Specificity of PCR products was confirmed by melting curve analyses. Prokaryotic SSU rRNA gene copy numbers were calculated as gene copy number g^−1^ of fresh material.

### 5.7 Ecological, phylogenetic, statistical and bioinformatic analysis

Amplification primers were removed from raw fastq files using Cutadapt v.2.0 (Martin, 2011). Amplicon sequence variants (ASVs) were inferred from quality filtered reads utilizing DADA2 v.1.10 (Callahan et al., 2016). Paired reads were merged, and reads between 251 and 255 bp length were conserved. Chimeras were removed using the removeBimeraDenovo function from the DADA2 package. Taxonomy was assigned utilizing the RDP Naive Bayesian Classifier (Wang et al., 2007) against the SILVA SSU rRNA reference alignment (Release 132, Quast et al., 2013). Sequences classified as chloroplast, mitochondrial, eukaryotic, or that did not match any taxonomic phylum were excluded from the dataset. Reads were filtered to remove ASVs that appeared in less than 5% of the samples and/or had less than 10 total counts. A total of 32,740 unique ASVs were aligned to the SILVA SSU rRNA reference alignment (Release 132, Quast et al., 2013) in mothur v.1.43 (Schloss et al., 2009), and an approximately maximum-likelihood tree was constructed using FastTree v.2.1 (Price et al., 2010). Finally, 10,068,980 high quality SSU rRNA sequence reads with a median depth of 49,619 reads per sample were used for subsequent analysis.

Shannon diversity index was estimated using the phyloseq v.1.26 package (McMurdie and Holmes, 2013). Non-metric multidimensional scaling (nMDS) ordination utilizing the Bray-Curtis dissimilarity distance was performed. Multivariate variation of the Bray-Curtis dissimilarity matrix was partitioned to microbiome compartment (bulk sediment, rhizosphere, and root), *S. alterniflora* phenotype (tall, medium, and short), depth (0-5 cm, and 5-10 cm), location (Sapelo Island, and Skidaway Island), and year (2018, and 2019) based on a PERMANOVA analysis with 999 permutations performed in vegan v. 2.5 (Oksanen et al. 2013). PERMANOVA analysis was run for the complete dataset and in subsets per microbiome compartment. Differential abundance analysis was performed to assess genera that were significantly enriched in specific plant compartments, and in zones of the marsh associated with different *S. alterniflora* phenotypes, using DESeq2 v.1.26 (Love et al., 2014).

To evaluate phylogenetic community structure within (alpha) and between (beta) communities, we quantified the nearest taxon index (NTI) and the beta nearest taxon index (βNTI), respectively (Stegen et al., 2012, 2013). NTI and βNTI indices were calculated as the number of standard deviations of the observed mean-nearest-taxon-distance (MNTD) and βMNTD from a null distribution (999 randomizations of all ASVs names across phylogenetic tree tips) using the picante package v. 1.8 (Kembel et al., 2010). For within community analysis, an NTI greater than +2 indicates that coexisting taxa are more closely related than expected by chance (phylogenetic clustering due to environmental filtering); while an NTI less than −2 indicates that coexisting taxa are more distantly related than expected by chance (phylogenetic overdispersion due to greater competition between closely related ASVs) (Stegen et al., 2012). For βNTI, we assessed pairwise comparisons for samples from the same plant compartment, *S. alterniflora* phenotype, and sampling event in order to evaluate if phylogenetic structure within communities (NTI) replicated at a greater scale (βNTI between samples occupying the same marsh microenvironment). A βNTI value <−2 or >+2 indicates less or greater than expected phylogenetic turnover between two samples than expected by chance, respectively (Stegen et. al., 2012).

The *S. alterniflora* core root microbiome was investigated. For this study, an ASV prevalence threshold was operationally defined by plotting the relative abundance and richness of the rhizosphere and root core microbiomes at 10% intervals from 0% to 100% ASV prevalence cutoffs (Fig. S8). A conservative prevalence cutoff of 60% was determined by visually inspecting a threshold in which richness remained stable at increasing cutoff values (Fig. S8). Finally, putative nitrifying, S oxidizing, S/sulfate reducing, and Fe oxidizing function was inferred based on homology of ASVs at the genus level with previously described prokaryotic species (Table S1).

## DATA AVAILABILITY

Sequence data is available in the BioProject database (http://ncbi.nlm.nih.gov/bioproject) under accessions PRJNA666636. Associated data and metadata, and R script for bioinformatic pipeline used in this study is available in https://github.com/kostka-lab/Spartina_GA_Core_Microbiome

## COMPETING INTERESTS

The authors declare that they have no competing interests

## AUTHORS CONTRIBUTIONS

J.L.R., M.K. and J.E.K conceived of the study; J.L.R., M.K., T.S., J.E.K collected samples from the field; J.L.R., and M.K. performed the experiment, and the data analyses. J.L.R., M.K. and J.E.K., wrote the manuscript; J.L.R., M.K. T.S., and J.E.K., provided valuable insight and ideas during numerous sessions of discussion. All authors provided critical comments on the manuscript and gave final approval for publication.

## FUNDING

This work was supported in part by an institutional grant (NA18OAR4170084) to the Georgia Sea Grant College Program from the National Sea Grant Office, National Oceanic and Atmospheric Administration, U.S. Department of Commerce and by a grant from the National Science Foundation (DEB 1754756). Any opinions, findings and conclusions or recommendations expressed in this material are those of the authors and do not necessarily reflect the views of the National Science Foundation.

## ACKNOWLEDGMENTS

Authors would like to acknowledge Christina Stoner and Jack Cenatempo, as well as teachers from the GCE Schoolyard Program for their field work assistance. This is contribution 1087 of the University of Georgia Marine Institute.

## Notes

### Competing Interest Statement

The authors have declared no competing interest.

## REFERENCES

1. An, T.T., Picardal, F.W., 2014. *Desulfocarbo indianensis* gen. nov., sp. nov., a benzoate-oxidizing, sulfate-reducing bacterium isolated from water extracted from a coal bed. International journal of systematic and evolutionary microbiology, 64:2907–2914.

2. Bahr, M., Crump, B.C., Klepac-Ceraj, V., Teske, A., Sogin, M.L., Hobbie, J.E., 2005. Molecular characterization of sulfate-reducing bacteria in a New England salt marsh. Environmental Microbiology, 7:1175–1185.

3. Barbier, E.B., Hacker, S.D., Kennedy, C., Koch, E.W., Stier, A.C., Silliman, B.R., 2011. The value of estuarine and coastal ecosystem services. Ecological Monographs, 81:169–193.

4. Barbieri, E., Ceccaroli, P., Saltarelli, R., Guidi, C., Potenza, L., Basaglia, M., et al., 2010. New evidence for nitrogen fixation within the Italian white truffle Tuber magnatum. Fungal biology, 114:936–942.

5. Beckler, J.S., Nuzzio, D.B., Taillefert, M., 2014. Development of single-step liquid chromatography methods with ultraviolet detection for the measurement of inorganic anions in marine waters. Limnology and Oceanography: Methods 12:563–576.

6. Bradley, P.M., Morris, J.T., 1990. Influence of oxygen and sulfide concentration on nitrogen uptake kinetics in *Spartina alterniflora*. Ecology, 71:282–287.

7. Brown, M. M., Friez, M. J., Lovell, C. R., 2003. Expression of nifH genes by diazotrophic bacteria in the rhizosphere of short form *Spartina alterniflora*. FEMS Microbiology Ecology, 43:411–417.

8. Bulgarelli, D., Schlaeppi, K., Spaepen, S., Van Themaat, E.V.L., Schulze-Lefert, P., 2013. Structure and functions of the bacterial microbiota of plants. Annual review of plant biology, 64:807–838.

9. Callahan, B.J., McMurdie, P.J., Rosen, M.J., Han, A.W., Johnson, A.J., Holmes, S.P., 2016. DADA2: High-resolution sample inference from Illumina amplicon data. Nature methods 13:581–583.

10. Caporaso, G., Lauber, C.L., Walters, W.A., Berg-Lyons, D., Lozupone, C.A., Turnbaugh, P.J., Fierer, N., Knight, R., 2011. Global patterns of 16S rRNA diversity at a depth of millions of sequences per sample. PNAS 108:4516–4522.

11. Carlson, P.R., Forrest, J., 1982. Uptake of dissolved sulfide by *Spartina alterniflora*: Evidence from natural sulfur isotope abundance ratios. Science, 216:633–635.

12. Cline, J.D., 1969. Spectrophotometric determination of hydrogen sulfide in natural waters. Limnol. Oceanogr. 14:454–458.

13. Craine, J.M., Brookshire, E.N.J., Cramer, M.D., Hasselquist, N.J., Koba, K., Marin-Spiotta, E., Wang, L., 2015. Ecological interpretations of nitrogen isotope ratios of terrestrial plants and soils. Plant and Soil, 396:1–26.

14. Crump, B.C., Wojahn, J.M., Tomas, F., Mueller, R.S., 2018. Metatranscriptomics and Amplicon Sequencing Reveal Mutualisms in Seagrass Microbiomes. Frontiers in Microbiology, 9:388.

15. Dai, J., Sun, M.-Y., Culp, R.A., Noakes, J.E., 2005. Changes in chemical and isotopic signatures of plant materials during degradation: Implication for assessing various organic inputs in estuarine systems. Geophysical Research Letters, 32: L13608.

16. Davis, D. A., Gamble, M. D., Bagwell, C. E., Bergholz, P. W., Lovell, C. R., 2011. Responses of salt marsh plant rhizosphere diazotroph assemblages to changes in marsh elevation, edaphic conditions and plant host species. Microbial ecology, 61:386–398.

17. Dini-Andreote, F., Stegen, J.C., van Elsas, J.D., Salles, J.F., 2015. Disentangling mechanisms that mediate the balance between stochastic and deterministic processes in microbial succession. Proceedings of the National Academy of Sciences, 112:E1326–E1332.

18. Dollhopf, S.L., Hyun, J.H., Smith, A.C., Adams, H.J., O’Brien, S., Kostka, J.E., 2005. Quantification of ammonia-oxidizing bacteria and factors controlling nitrification in salt marsh sediments. Applied and Environmental Microbiology, 71:240–246.

19. Dubilier, N., Bergin, C., Lott, C., 2008. Symbiotic diversity in marine animals: the art of harnessing chemosynthesis. Nature Reviews Microbiology, 6:725–740.

20. Edwards, J., Johnson, C., Santos-Medellín, C., Lurie, E., Podishetty, N.K., Bhatnagar, S., Eisen, J.A., Sundaresan, V., 2015. Structure, variation, and assembly of the root-associated microbiomes of rice. Proceedings of the National Academy of Sciences, 112:E911–E920.

21. Ember, L.M., Williams, D.F., Morris, J.T., 1987. Processes that influence carbon isotope variations in salt marsh sediments. Marine Ecology Progress Series, 36:33–42.

22. Fogel, M. L., Sprague, E. K., Gize, A. P., Frey, R. W., 1989. Diagenesis of organic matter in Georgia salt marshes. Estuarine, Coastal and Shelf Science, 28:211–230.

23. Freedman, Z., Zak, D.R., 2015. Soil bacterial communities are shaped by temporal and environmental filtering: evidence from a long-term chronosequence. Environmental microbiology, 17:3208–3218.

24. Fukami, T., 2015. Historical contingency in community assembly: integrating niches, species pools, and priority effects. Annual Review of Ecology, Evolution, and Systematics, 46:1–23.

25. Gagnon, K., Rinde, E., Bengil, E.G., Carugati, L., Christianen, M.J., Danovaro, R., Gambi, C., Govers, L.L., Kipson, S., Meysick, L., Pajusalu, L., 2020. Facilitating foundation species: The potential for plant–bivalve interactions to improve habitat restoration success. Journal of Applied Ecology. DOI: 10.1111/1365-2664.13605

26. Gandy, E.L., Yoch, D.C., 1988. relationship between nitrogen-fixing sulfate reducers and fermenters in salt marsh sediments and roots of *Spartina alterniflora*. Applied and Environmental Microbiology, 54:2031–2036.

27. Garcia-Robledo, E., Corzo, A., Papaspyrou, S., 2014. A fast and direct spectrophotometric method for the sequential determination of nitrate and nitrite at low concentrations in small volumes. Marine Chemistry, 162:30–36.

28. Gebrehiwet, T., Koretsky, C.M., Krishnamurthy, R.V., 2008. Influence of *Spartina* and *Juncus* on saltmarsh sediments. III. Organic geochemistry. Chemical Geology, 255:114–119.

29. Giurgevich, J.R., Dunn, E.L., 1979. Seasonal Patterns of CO2 and Water Vapor Exchange of the Tall and Short Height Forms of *Spartina alterniflora* Loisel in a Georgia Salt Marsh. Oecologia, 43:139–156.

30. Gribsholt, B., Kostka, J. E., & Kristensen, E., 2003. Impact of fiddler crabs and plant roots on sediment biogeochemistry in a Georgia saltmarsh. Marine Ecology Progress Series, 259:237–251.

31. Hassani, M.A., Durán, P., Hacquard, S., 2018. Microbial interactions within the plant holobiont. Microbiome, 6:58.

32. Higashioka, Y., Kojima, H., Watanabe, M., Fukui, M., 2013. *Desulfatitalea tepidiphila* gen. nov., sp. nov., a sulfate-reducing bacterium isolated from tidal flat sediment. International journal of systematic and evolutionary microbiology, 63:761–765.

33. Higashioka, Y., Kojima, H., Watanabe, T., & Fukui, M., 2015. Draft genome sequence of *Desulfatitalea tepidiphila* S28bFT. Genome announcements, 3(6), e01326–15.

34. Hong, Y., Liao, D., Hu, A., Wang, H., Chen, J., Khan, S., Su, J., Li, H., 2015. Diversity of endophytic and rhizoplane bacterial communities associated with exotic *Spartina alterniflora* and native mangrove using Illumina amplicon sequencing. Canadian journal of microbiology, 61:723–733.

35. Hopkinson, C.S., Wolanski, E., Brinson, M.M., Cahoon, D.R., and Perillo, G.M.E., 2019. Coastal Wetlands: A Synthesis. In: G.M.E., Perillo, E., Wolanski, D.R., Cahoon, and C.S., Hopkinson. (Eds.) Coastal Wetlands, Second Edition: An Integrated and Ecosystem Approach. Elsevier, pp. 1–75.

36. Howarth, R.W., 1984. The ecological significance of sulfur in the energy dynamics of salt marsh and coastal marine sediments. Biogeochemistry, 1:5–27.

37. Howes, B.L., Dacey, J.W.H., Goehringer, D.D., 1986. Factors controlling the growth form of *Spartina alterniflora*: feedbacks between above-ground production, sediment oxidation, nitrogen and salinity. Journal of Ecology, 74:881–898.

38. Hwang, Y-S., Morris, J.T., 1994. Whole-plant gas exchange responses of *Spartina alterniflora* (Poaceae) to a range of constant and transient salinities. American Journal of Botany, 81:659–665.

39. Hyun, J.H., Smith, A.C., Kostka, J.E., 2007. Relative contributions of sulfate-and iron (III) reduction to organic matter mineralization and process controls in contrasting habitats of the Georgia saltmarsh. Applied Geochemistry, 22:2637–2651.

40. Janssen, B., 1996. Nitrogen mineralization in relation to C:N ratio and decomposability of organic materials. Plant Soil. 181:39–45

41. Joye, S.B., Hollibaugh, J.T., 1995. influence of sulfide inhibition of nitrification on nitrogen regeneration in sediments. Science, 270:623–625.

42. Kembel, S., Cowan, P., Helmus, M., Cornwell, W., Morlon, H., Ackerly, D., Blomberg, S., Webb, C., 2010. Picante: R tools for integrating phylogenies and ecology. Bioinformatics, 26:1463–1464.

43. Kirwan, M.L., Guntenspergen, G.R., Morris, J.T., 2009. Latitudinal trends in *Spartina alterniflora* productivity and the response of coastal marshes to global change. Global Change Biology, 15:1982–1989.

44. Koch, M.S., Mendelssohn, I.A., McKee, K.L., 1990. Mechanism for the hydrogen sulfide-induced growth limitation in wetland macrophytes. Limnology and Oceanography, 35:399–408.

45. Kolton, M., Rolando, J.L., Kostka, J.E., 2020. Elucidation of the rhizosphere microbiome linked to *Spartina alterniflora* phenotype in a salt marsh on Skidaway Island, Georgia, USA. FEMS Microbiology Ecology, 96:fiaa026.

46. Koop-Jakobsen, K., Mueller, P., Meier, R.J., Liebsch, G., Jensen, K., 2018. Plant-sediment interactions in salt marshes–an optode imaging study of O2, pH, and CO2 gradients in the rhizosphere. Frontiers in plant science, 9:541.

47. Koretsky, C.M., Moore, C.M., Lowe, K.L., Meile, C., DiChristina, T.J., Van Cappellen, P., 2003. Seasonal oscillation of microbial iron and sulfate reduction in saltmarsh sediments (Sapelo Island, GA, USA). Biogeochemistry, 64:179–203.

48. Kostka, J.E., Gribsholt, B., Petrie, E., Dalton, D., Skelton, H.,Kristensen, E., 2002a. The rates and pathways of carbon oxidation in bioturbated saltmarsh sediments. Limnology and Oceanography, 47:230–240.

49. Kostka, J.E., Roychoudhury, A., Van Cappellen, P., 2002b. Rates and controls of anaerobic microbial respiration across spatial and temporal gradients in saltmarsh sediments. Biogeochemistry, 60:49–76.

50. Lee, R.W., Kraus, D.W., Doeller, J.E., 1999. Oxidation of sulfide by *Spartina alterniflora* roots. Limnology and Oceanography, 44:1155–1159.

51. Lever, M.A., Torti, A., Eickenbusch, P., Michaud, A.B., Šantl-Temkiv, T., Jørgensen, B.B., 2015. A modular method for the extraction of DNA and RNA, and the separation of DNA pools from diverse environmental sample types. Frontiers in Microbiology, 6:476.

52. Lim, S.J., Davis, B.G., Gill, D.E., Walton, J., Nachman, E., Engel, A.S., Anderson, L.C. and Campbell, B.J., 2019a. Taxonomic and functional heterogeneity of the gill microbiome in a symbiotic coastal mangrove lucinid species. The ISME journal, 13:902–920.

53. Lim, S.J., Alexander, L., Engel, A.S., Paterson, A.T., Anderson, L.C., Campbell, B.J., 2019b. Extensive thioautotrophic gill endosymbiont diversity within a single *Ctena orbiculate* (Bivalvia: Lucinidae) population and implications for defining host-symbiont specificity and species recognition. mSystems, 4:e00280–19.

54. Lin, L., Liu, W., Zhang, M., Lin, X., Zhang, Y., Tian, Y., 2019. Different height forms of *Spartina alterniflora* might select their own rhizospheric bacterial communities in southern coast of China. Microbial Ecology, 77:124–135.

55. Liu, Y., Zhu, A., Tan, H., Cao, L., Zhang, R., 2019. Engineering banana endosphere microbiome to improve Fusarium wilt resistance in banana. Microbiome, 7:74.

56. Love, M.I., Huber, W., Anders, S., 2014. Moderated estimation of fold change and dispersion for RNA-seq data with DESeq2. Genome Biology, 15:550.

57. Lovell, C.R., Piceno, Y.M., Quattro, J.M., Bagwell, C.E., 2000. Molecular analysis of diazotroph diversity in the rhizosphere of the smooth cordgrass, *Spartina alterniflora*. Applied and Environmental Microbiology, 66:3814–3822.

58. Lundberg, D.S., Yourstone, S., Mieczkowski, P., Jones, C.D., Dangl, J.L., 2013. Practical innovations for high-throughput amplicon sequencing. Nature Methods, 10:999–1002.

59. Maricle, B.R., Lee, R.W., 2002. Aerenchyma development and oxygen transport in the estuarine cordgrasses Spartina alterniflora and S. anglica. Aquatic Botany, 74:109–120.

60. Martin, M., 2011. Cutadapt removes adapter sequences from high-throughput sequencing reads. EMBnet.Journal, 17:10–12.

61. Martin, B.C., Alarcon, M.S., Gleeson, D., Middleton, J.A., Fraser, M.W., Ryan, M.H., Holmer, M., Kendrick, G.A., Kilminster, K., 2020a. Root microbiomes as indicators of seagrass health. FEMS Microbiology Ecology, 96:fiz201.

62. Martin, B.C., Middleton, J.A., Fraser, M.W., Marshall, I.P., Scholz, V.V. and Schmidt, H., 2020b. Cutting out the middle clam: lucinid endosymbiotic bacteria are also associated with seagrass roots worldwide. The ISME journal, 14:2901–2905.

63. McMurdie, P.J., Holmes, S., 2013. phyloseq: an R package for reproducible interactive analysis and graphics of microbiome census data. PloS one, 8(4).

64. Mcowen, C.J., Weatherdon, L.V., Van Bochove, J.W., Sullivan, E., Blyth, S., Zockler, C., Stanwell-Smith, D., Kingston, N., Martin, C.S., Spalding, M., Fletcher, S., 2017. A global map of saltmarshes. Biodiversity data journal, 5: e11764.

65. Meier, D. V., Pjevac, P., Bach, W., Hourdez, S., Girguis, P. R., Vidoudez, C., et al., 2017. Niche partitioning of diverse sulfur-oxidizing bacteria at hydrothermal vents. The ISME Journal, 11:1545–1558.

66. Mendelssohn, I.A., 1979. the influence of nitrogen level, form, and application method on the growth response of *Spartina alterniflora* in North Carolina. Estuaries, 2:106–112.

67. Mendelssohn, I.A., McKee, K.L., 1992. Indicators of environmental stress in wetland plants. In Ecological indicators (pp. 603–624). Springer, Boston, MA.

68. Mendelssohn, I.A., Morris, J.T., 2000. Eco-physiological controls on the productivity of *Spartina alterniflora* Loisel. In: Weinstein, M.P., Kreeger, D.A. (Eds.). Concepts and controversies in tidal marsh ecology. Kluwer Academic Publishers, pp. 59–80.

69. Mitsch, W. J., Gosselink, J. G., 1993. Wetlands. Van Nostrand Reinhold, New York, New York, USA.

70. Morris, J.T., Dacey, J.W.H., 1984. Effects of O2 on ammonium uptake and root respiration by *Spartina alterniflora*. American Journal of Botany, 71:979–985.

71. Morris, J.T., Haley, C., Krest, R., 1996. Effects of sulfide concentrations on growth and dimethylsulphoniopropionate (DMSP) concentration in *Spartina alterniflora*. In: Kiene, R., Visscher, R., Keller, M., Kirst, G. (Eds.) Biological and environmental chemistry of DMSP and related sulfonium compounds. Plenum, pp. 87–95

72. Moulana, A., Anderson, R. E., Fortunato, C. S., Huber, J. A., 2020. Selection is a significant driver of gene gain and loss in the pangenome of the bacterial genus *Sulfurovum* in geographically distinct deep-sea hydrothermal vents. Msystems, 5:e00673–19.

73. Murphy, A.E., Bulseco, A.N., Ackerman, R., Vineis, J.H. and Bowen, J.L., 2020. Sulphide addition favours respiratory ammonification (DNRA) over complete denitrification and alters the active microbial community in salt marsh sediments. Environmental Microbiology, 22:2124–2139.

74. Oksanen, J., Blanchet, F.G., Kindt, R., Legendre, P., Minchin, P.R., O’hara, R.B., Simpson, G.L., Solymos, P., Stevens, M.H.H., Wagner, H., Oksanen, M.J., 2013. Package ‘vegan’. Community ecology package, version, 2(9), pp.1–295.

75. Petersen, J.M., Kemper, A., Gruber-Vodicka, H., Cardini, U., Van Der Geest, M., Kleiner, M., Bulgheresi, S., Mußmann, M., Herbold, C., Seah, B.K., Antony, C.P., 2017. Chemosynthetic symbionts of marine invertebrate animals are capable of nitrogen fixation. Nature microbiology, 2:16195.

76. Peterson, B.J., 1999. Stable isotopes as tracers of organic matter input and transfer in benthic food webs: A review. Acta Oecologica, 20:479–487.

77. Price, M.N., Dehal, P.S., Arkin, A.P., 2010. FastTree 2 – approximately maximum-likelihood trees for large alignments. PLoS ONE, 5:e9490.

78. Quast, C., Pruesse, E., Yilmaz, P., Gerken, J., Schweer, T., Yarza, P., Peplies, J., Glöckner, F.O., 2013. The SILVA ribosomal RNA gene database project: improved data processing and web-based tools. Nucledic Acids Research, 41:D590–D596.

79. Ramírez, D.A., Yactayo, W., Rens, L.R., Rolando, J.L., Palacios, S., De Mendiburu, F., Mares, V., Barreda, C., Loayza, H., Monneveux, P., Zotarelli, L., Khan, A., Quiroz, R., 2016. Defining biological thresholds associated to plant water status for monitoring water restriction effects: Stomatal conductance and photosynthesis recovery as key indicators in potato. Agricultural Water Management, 177:369–378.

80. Ramnarine, R., Voroney, R.P., Wagner-Riddle, C., Dunfield, K.E., 2011. Carbonate removal by acid fumigation for measuring the δ^13^C of soil organic carbon. Canadian Journal of Soil Science, 91:247–250.

81. Simmons, T., Caddell, D.F., Deng, S., Coleman-Derr, D., 2018. Exploring the root microbiome: extracting bacterial community data from the soil, rhizosphere, and root endosphere. Journal of Visualized Experiments, 135:e57561.

82. Schloss, P.D., Westcott, S.L., Ryabin, T., Hall, J.R., Hartmann, M., Hollister, E.B., Lesniewski, R.A., Oakley, B.B., Parks, D.H., Robinson, C.J., Sahl, J.W., 2009. Introducing mothur: open-source, platform-independent, community-supported software for describing and comparing microbial communities. Applied and Environmental Microbiology, 75:7537–7541.

83. Shade, A., Handelsman, J., 2012. Beyond the Venn diagram: the hunt for a core microbiome. Environmental microbiology, 14:4–12.

84. Sims, G.K., Ellsworth, T.R., Mulvaney, R.L., 1995. Microscale determination of inorganic nitrogen in water and soil extracts. Communications in Soil Science and Plant Analysis, 26:303–316.

85. Spivak, A.C., Reeve, J., 2015. Rapid cycling of recently fixed carbon in a *Spartina alterniflora* system: a stable isotope tracer experiment. Biogeochemistry 125:97–114.

86. Stegen, J.C., Lin, X., Konopka, A.E., Fredrickson, J.K., 2012. Stochastic and deterministic assembly processes in subsurface microbial communities. The ISME journal, 6:1653–1664.

87. Stegen, J.C., Lin, X., Fredrickson, J.K., Chen, X., Kennedy, D.W., Murray, C.J., Rockhold, M.L., Konopka, A., 2013. Quantifying community assembly processes and identifying features that impose them. The ISME journal, 7:2069–2079.

88. Stookey, L.L., 1970. Ferrozine – a new spectrophotometric reagent for iron. Anal. Chem. 43:779–781.

89. Suzuki, Y., Kojima, S., Sasaki, T., Suzuki, M., Utsumi, T., Watanabe, H., et al., 2006. Host-symbiont relationships in hydrothermal vent gastropods of the genus Alviniconcha from the Southwest Pacific. Applied and Environmental Microbiology, 72:1388–1393.

90. Suzuki, D., Li, Z., Cui, X., Zhang, C., Katayama, A., 2014. Reclassification of *Desulfobacterium anilini* as *Desulfatiglans anilini* comb. nov. within *Desulfatiglans* gen. nov., and description of a 4-chlorophenol-degrading sulfate-reducing bacterium, *Desulfatiglans parachlorophenolica* sp. nov. International journal of systematic and evolutionary microbiology, 64:3081–3086.

91. Thajudeen, J., Yousuf, J., Veetil, V. P., Varghese, S., Singh, A., Abdulla, M. H., 2017. Nitrogen fixing bacterial diversity in a tropical estuarine sediments. World Journal of Microbiology and Biotechnology, 33:41.

92. Thomas, F., Giblin, A.E., Cardon, Z.G., Sievert, S.M., 2014. Rhizosphere heterogeneity shapes abundance and activity of sulfur-oxidizing bacteria in vegetated salt marsh sediments. Frontiers in microbiology, 5:309.

93. Tobias C.R., Neubauer S.C., 2019. Salt marsh biogeochemistry—an overview. In: Perillo, G.M.E., Wolanski, E., Cahoon D.R., Brinson, M.M., (Eds.). Coastal wetlands: an integrated ecological approach. Elsevier, pp. 445–492.

94. Toju, H., Peay, K.G., Yamamichi, M., Narisawa, K., Hiruma, K., Naito, K., Fukuda, S., Ushio, M., Nakaoka, S., Onoda, Y., Yoshida, K., 2018. Core microbiomes for sustainable agroecosystems. Nature Plants, 4:247–257.

95. Trivedi, P., Leach, J. E., Tringe, S. G., Sa, T., Singh, B. K., 2020. Plant–microbiome interactions: from community assembly to plant health. Nature Reviews Microbiology, 18:607–621.

96. Valiela, I., Teal, J.M., Deuser, W.G., 1978. The nature of growth forms in the salt marsh grass *Spartina alterniflora*. The American Naturalist 112:461–470

97. Wang, Q., Garrity, G.M., Tiedje, J.M., Cole, J.R., 2007. Naive Bayesian classifier for rapid assignment of rRNA sequences into the new bacterial taxonomy. Applied and environmental microbiology, 73:5261–5267.

98. Watanabe, F.S., Olsen, S.R., 1965. Test of an ascorbic acid method for determining phosphorus in water and NaHCO3 extracts from soil. Soil Science Society Proceedings, 29:677–678.

99. White, J.F., Kingsley, K.L., Zhang, Q., Verma, R., Obi, N., Dvinskikh, S., Elmore, M.T., Verma, S.K., Gond, S.K., Kowalski, K.P., 2019. Endophytic microbes and their potential applications in crop management. Pest management science, 75:2558–2565.

100. Whiting, G.J., Gandy, E.L., Yoch, D.C., 1986. Tight coupling of root-associated nitrogen fixation and plant photosynthesis in the salt marsh grass *Spartina alterniflora* and carbon dioxide enhancement of nitrogenase activity. Applied and Environmental Microbiology, 52:108–113.

101. Wieski, K., Pennings, S., 2014. Climate Drivers of *Spartina alterniflora* saltmarsh production in Georgia, USA. Ecosystems, 17:473–484.

102. Zhang, L., Zhang, W., Li, Q., Cui, R., Wang, Z., Wang, Y., Zhang, Y.Z., Ding, W., Shen, X., 2020. Deciphering the root endosphere microbiome of the desert plant *Alhagi sparsifolia* for drought resistance-promoting bacteria. Applied and Environmental Microbiology, 86:e02863–19.

103. Zheng, Y., Hou, L., Liu, M., Yin, G., Gao, J., Jiang, X., Lin, X., Li, X., Yu, C., Wang, R., 2016. Community composition and activity of anaerobic ammonium oxidation bacteria in the rhizosphere of salt-marsh grass *Spartina alterniflora*. Applied Microbiology and Biotechnology, 100:8203–8212.

104. Zogg, G.P., Travis, S.E., Brazeau, D.A., 2018. Strong associations between plant genotypes and bacterial communities in a natural salt marsh. Ecology and Evolution 8:4721–4730.

