## Supplementary Tables and Figures for "The core root microbiome of *Spartina alterniflora* is predominated by sulfur-oxidizing and sulfate-reducing bacteria in Georgia salt marshes, USA"

**Supplemental information**

**Table S2:** Allometric equations used to estimate *S. alterniflora* biomass by plant phenotype according to Wieski and Pennings (2014)

| ***S. alterniflora* phenotype** | **Equation** |
|---|---|
| Tall | $\ln(biomass) = -6.095 + 1.760 * \ln(height)$ |
| Medium and short | $\ln(biomass) = -6.934 + 1.973 * \ln(height)$ |

Biomass: Shoot dry mass (g)

Height: Shoot height (cm)

**Table S3**: Extracellular enzyme substrates employed in the study identified by Chemical Abstracts Service (CAS) and Enzyme Commission (EC) designations.

| **Exoenzyme** | **Substrate name** | **CAS number** | **EC number** |
|---|---|---|---|
| β-Glucosidase | 4-Methyllumbelliferyl β-D-glucopyranoside | 18997-57-4 | 3.2.1.21 |
| β-1,4-N-acetylglucosaminidase (chitinase) | 4-Methyllumbelliferyl N-acetyl-β-D-glucosaminide | 37067-30-4 | 3.2.1.14 |
| Phosphatase | 4-Methyllumbelliferyl phosphate | 3368-04-5 | 3.1.3.2 / 3.1.3.1 |

**Table S4:** PCR amplification conditions employed in the study.

| **Reaction** | **Primer set** | **Initial Denaturat.** | **Denaturat.** | **PNAs annealing** | **Primers annealing** | **Extension** | **Final extension** |
|---|---|---|---|---|---|---|---|
| 1st PCR | 515F/806R[1] | 2m, 95°C | 94°C, 45s | 78°C, 10s | 50°C, 60s | 72°C, 60s | 10m, 72°C |
| | | | x28 | | | | |
| 2nd PCR | Fluidigm 10-base barcodes | 5m, 95°C | 94°C, 30s | - | 60°C, 30s | 72°C, 30s | 5m, 72°C |
| | | | x8 | | | | |
| qPCR | 515F/806R[1] | 2m, 95°C | 94°C, 45s | 78°C, 10s | 50°C, 60s | 72°C, 90s | - |
| | | | x40 | | | | |

[1]Caporaso et al. (2011)

**Fig. S1.** Satellite images of study areas: (a) Sapelo Island GCE-6 site, and (b) Skidaway Island SERF site.

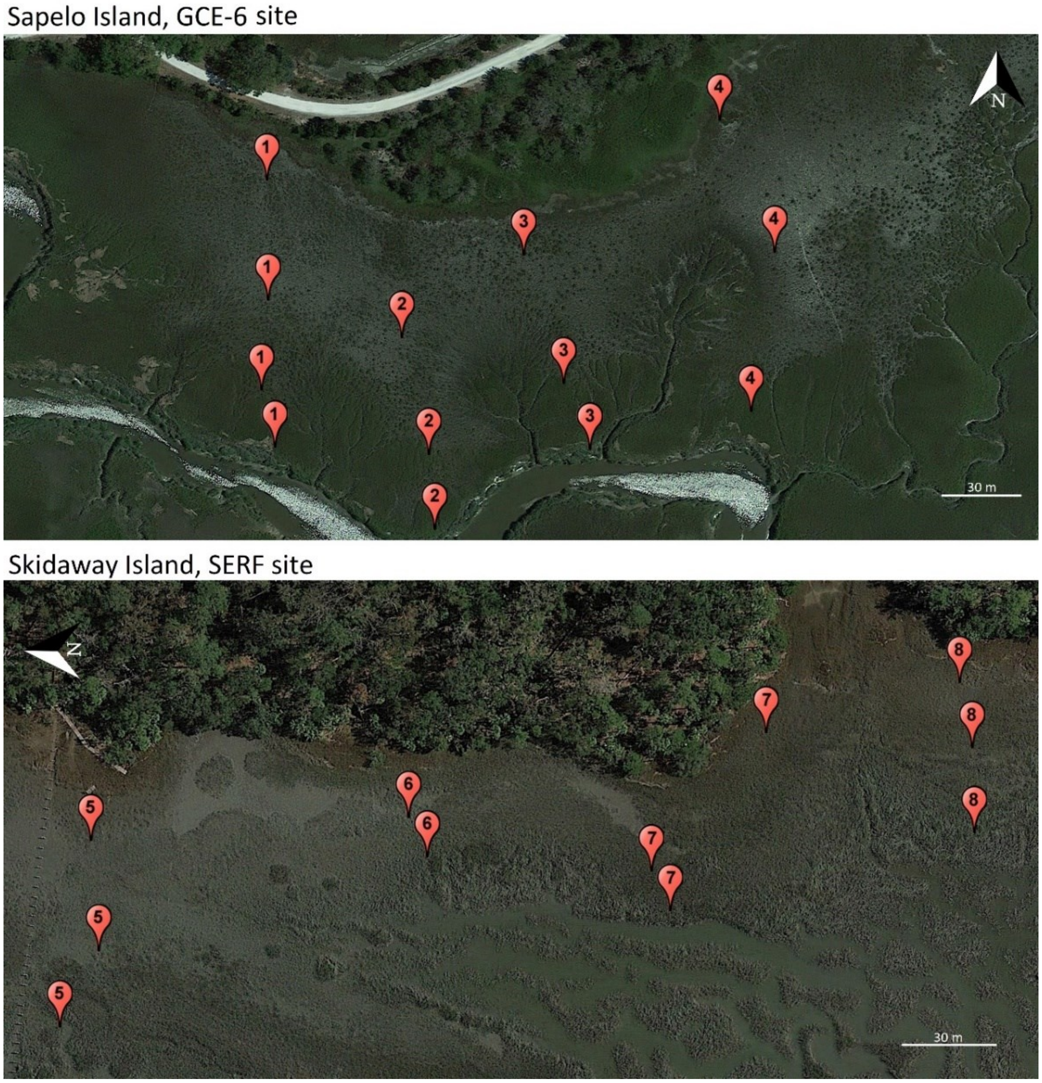

**Fig. S2.** Site ecological characterization. Boxplots of leaf N concentration, density of crab burrows, marsh periwinkle snail density, sediment redox potential (Eh), sediment pH, and porewater salinity grouped by *S. alterniflora* phenotype.

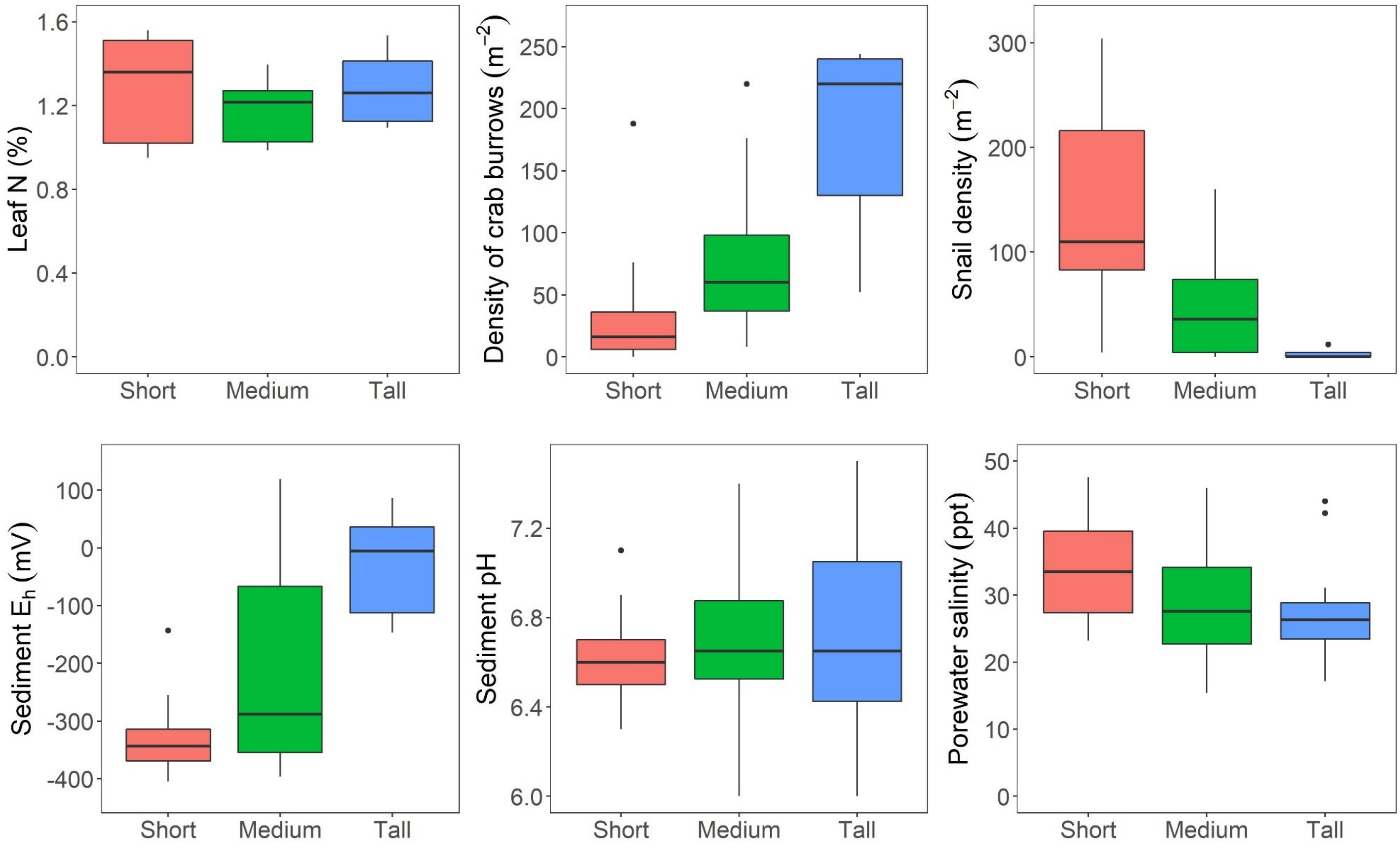

**Fig. S3.** Porewater chemistry characterization by *S. alterniflora* phenotype. Boxplots of ammonium, nitrate, phosphate, total sulfides, Fe(II), and Fe(III) porewater concentration per *S. alterniflora* phenotype.

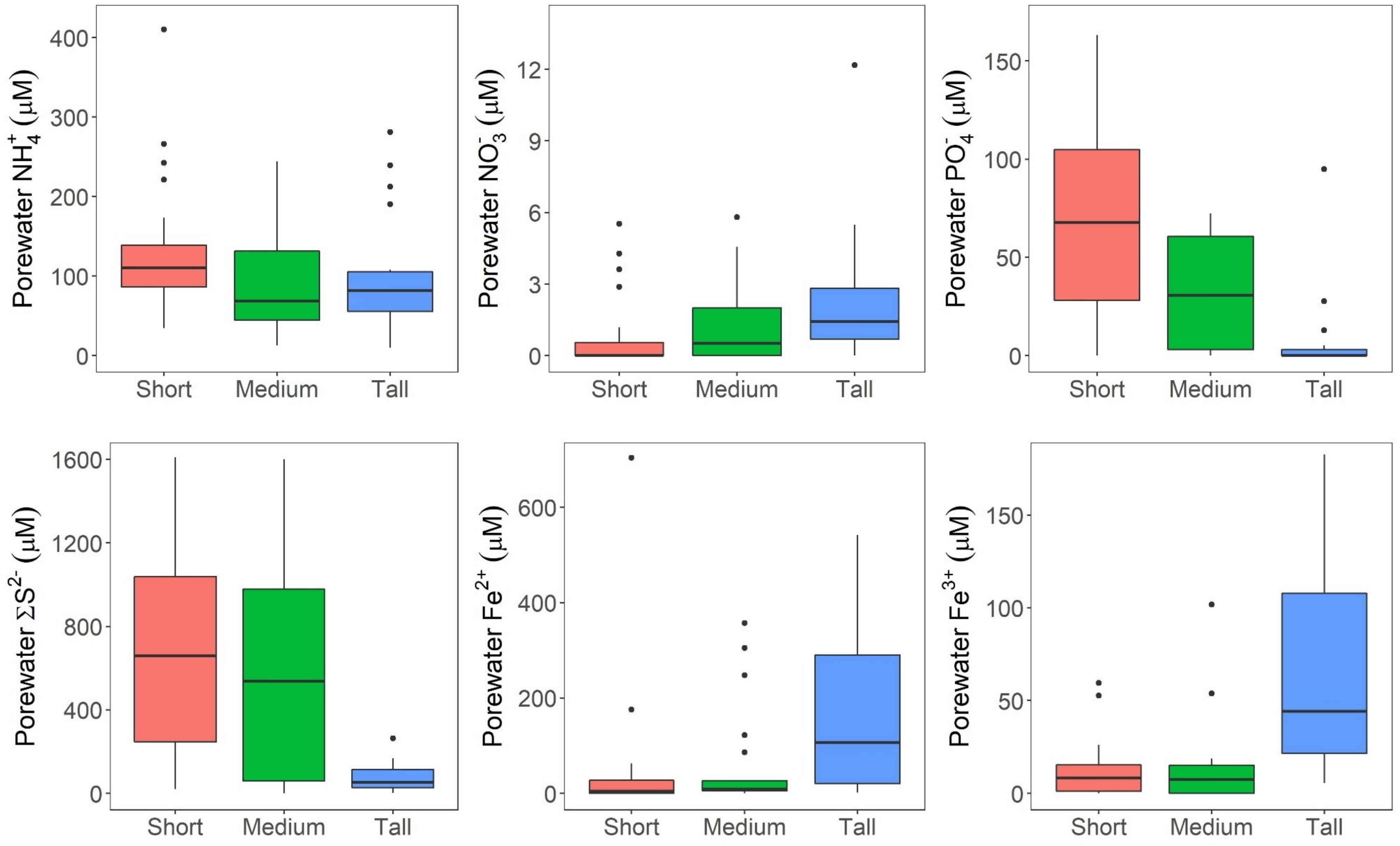

**Fig. S4.** C and N isotopic natural abundance by *S. alterniflora* phenotype. Boxplots of leaf $\delta^{15}N$, sediment $\delta^{15}N$, difference between leaf and sediment $\delta^{15}N$ ($\Delta^{15}N$), leaf $\delta^{13}C$, sediment $\delta^{13}C$, and difference between leaf and sediment $\delta^{13}C$ ($\Delta^{13}C$).

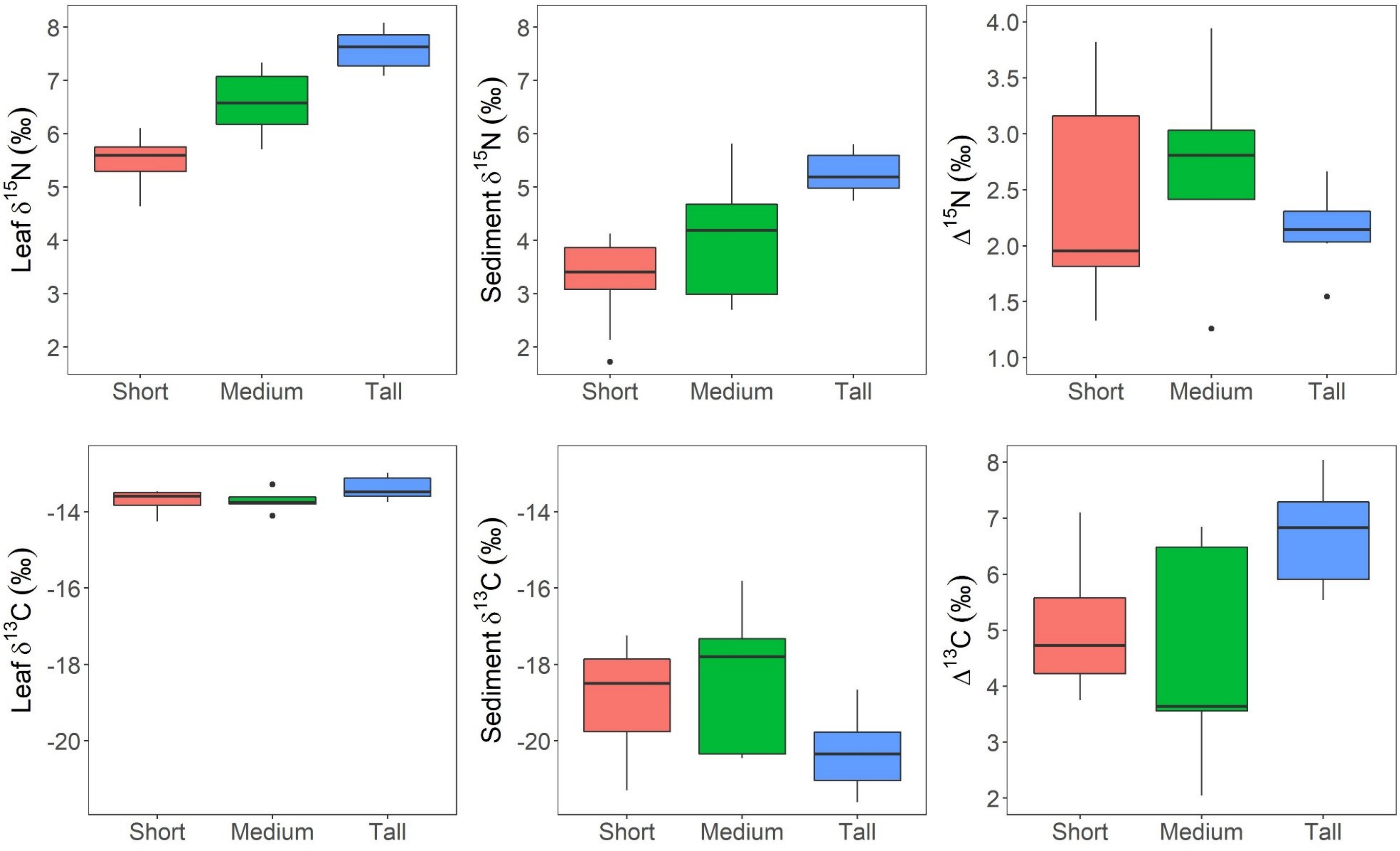

**Fig. S5.** Nearest taxon index (NTI) (a), and β-nearest taxon index (βNTI) (b) density plots per microbiome compartment. Dashed lines in -2 and, 2, represents interpretation thresholds. Continuous line represents median value according to index and microbiome compartment.

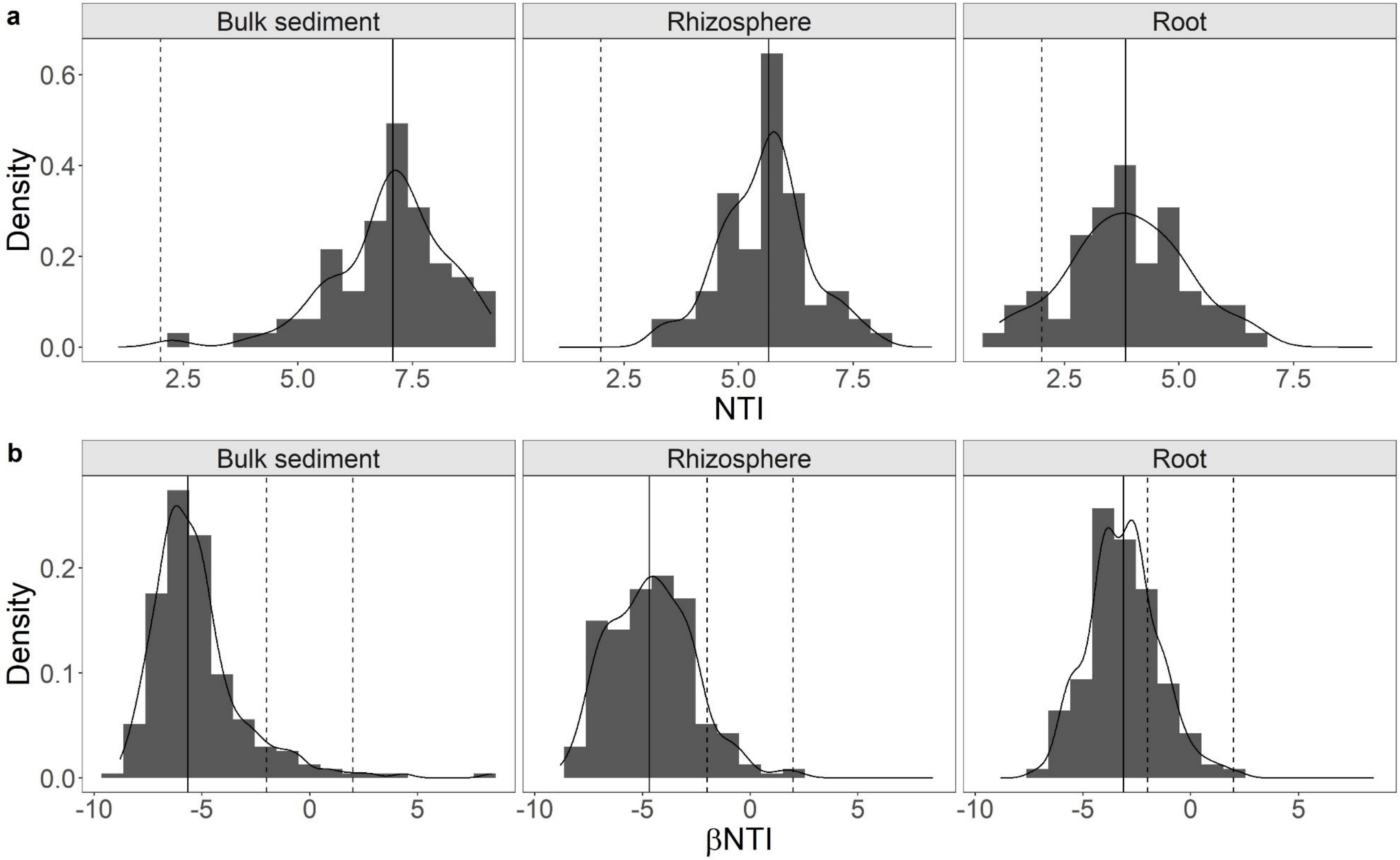

**Fig. S6.** Prokaryotic relative abundance partitioned at the phylum level according to microbiome compartment and *S. alterniflora* phenotype.

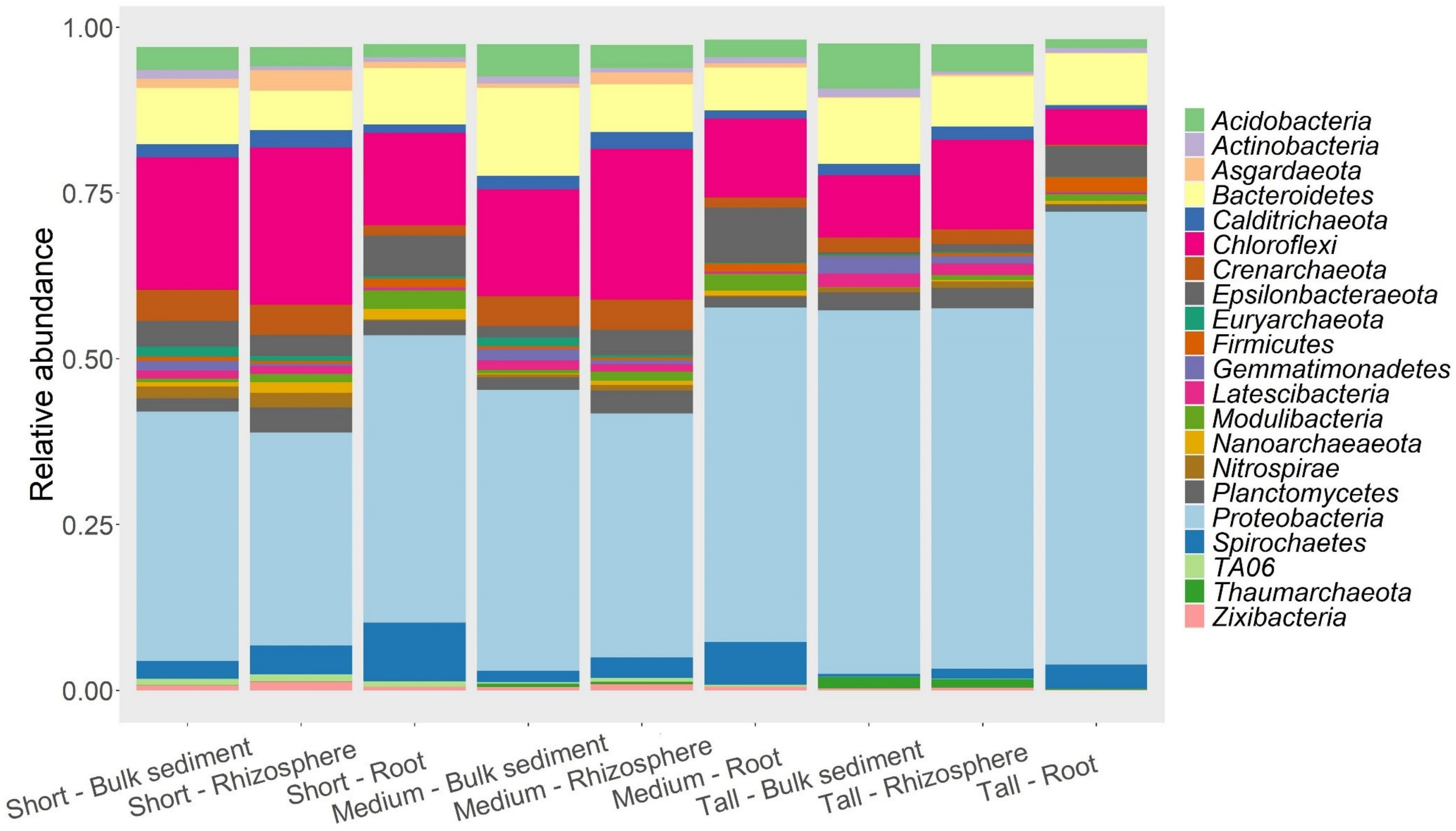

**Fig. S7.** Prokaryotic genera significantly enriched in the root in comparison to the bulk sediment compartments (a), and tall compared to short *S. alterniflora* phenotype (b) as assessed by DESeq2. Values represent the following: Yellow: Aerobic/facultative-anaerobic chemoheterotrophy, Green: N fixation, White: C fixation, Red: Nitrification, Blue: S oxidation, Black: S reduction, Brown: Methylotrophy, Purple: Metal reduction

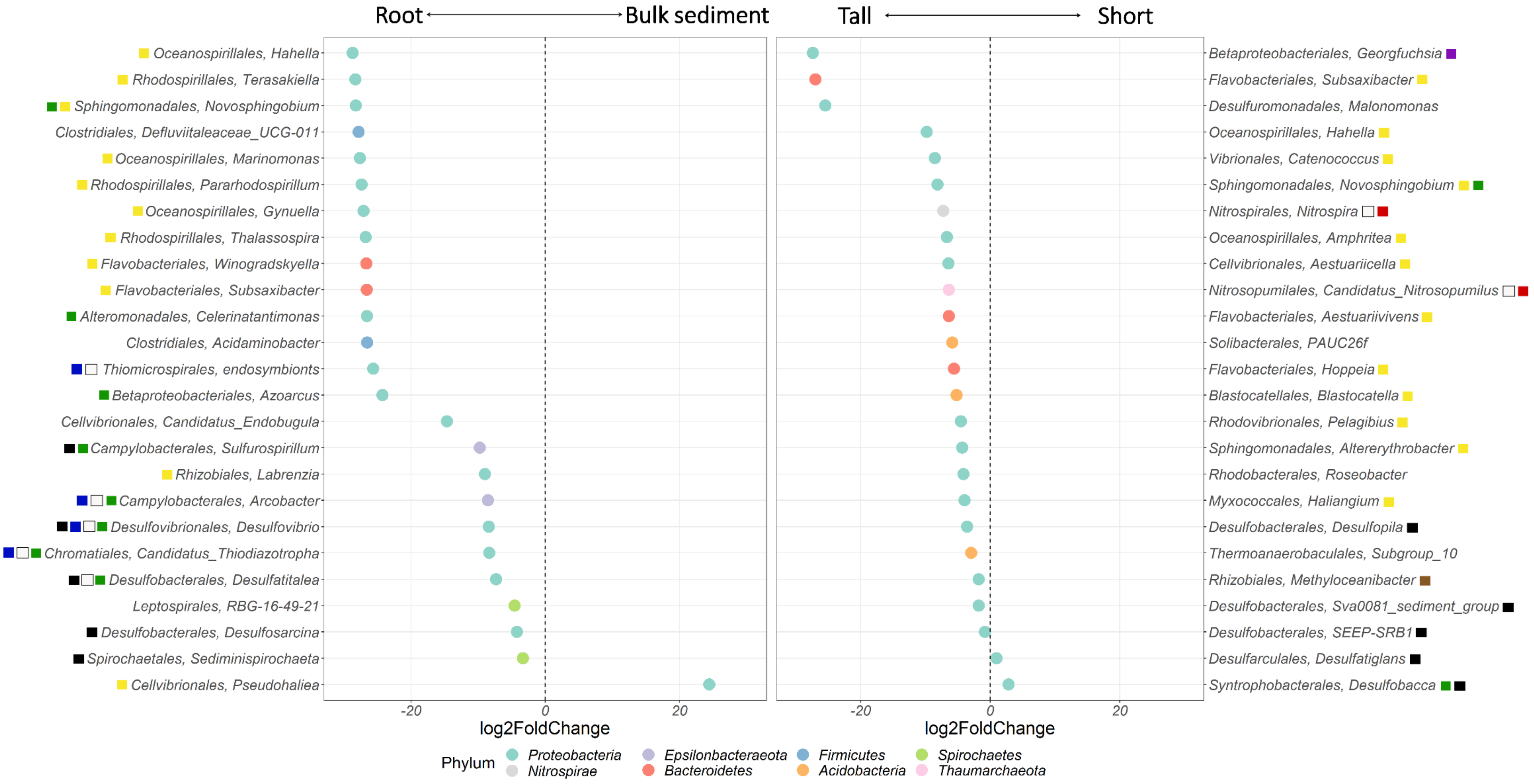

**Fig. S8.** Analysis of the core microbiome by accumulated richness and relative abundance with species prevalence cutoff thresholds at 10% intervals from 0% to 100% ASVs prevalence cutoffs.

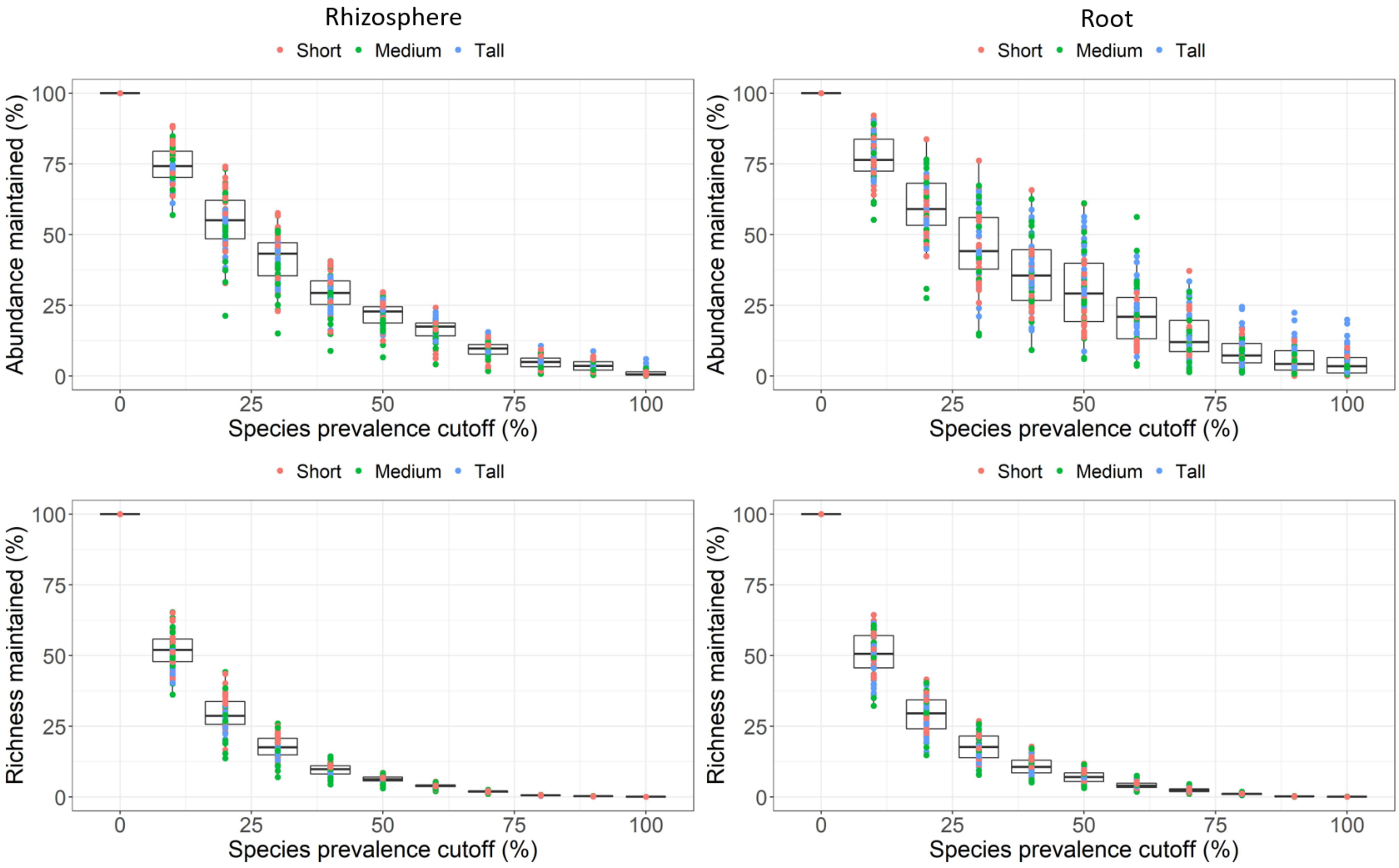
